# Coexistence in Periodic Environments

**DOI:** 10.1101/2023.02.24.529749

**Authors:** Alexa M Scott, Carling Bieg, Bailey C McMeans, Kevin S McCann

## Abstract

Climate change and other anthropogenic impacts are rapidly altering natural environmental periodicities on a variety of time scales. Despite this, a general theoretical foundation describing the role of periodic environmental variation in structuring species interactions and ecological communities is still underdeveloped. Alarmingly, this leaves us unprepared to understand and predict implications for the maintenance of biodiversity under global change. Here, we extend a two-species Lotka-Volterra competition model that incorporates periodic forcing between seasons of high and low production to investigate the effects of changing environmental patterns on species coexistence. Towards this, we define coexistence criteria for periodic environments by approximating isocline solutions akin to classical coexistence outcomes. This analytical approach illustrates that periodic environments (i.e., seasonality) in and of themselves can mediate different competitive outcomes, and these patterns are general across varying time scales. Importantly, species coexistence may be incredibly sensitive to changes in these abiotic periods, suggesting that climate change has the potential to drastically impact the maintenance of biodiversity in the future.

## Introduction

Nature is abundant with a diverse array of periodic climate signals (Jackson et al. 2021; Pokorný 2021). The complex variations in temperature over time (Jiang and Morin 2007; Klausmeier 2010), for example, can be decomposed into different lengths of underlying periodicities using spectral analysis, revealing a complex mosaic of short (e.g., seconds, minutes, hours, days), medium (e.g., months, years), long (e.g., multi-decadal), and very long natural periods (e.g., 100s to 1000s of years) (Forrest and Miller-Rushing 2010; Vasconcellos et al. 2011; Huntly et al. 2021; Joseph and Kumar 2021; Pokorný 2021). The regularity of these environmental periods allows for species to adapt and respond to them (Bernhardt et al. 2020; Fretwell 1972; Shuter et al. 2012; Tonkin et al. 2017), meaning that nature has evolved around, and within, these complex temporal abiotic signatures (Mathias and Chesson 2013; Varpe 2017; Rudolf 2019). Despite the long-known recognition of nature’s complex abiotic palette (White and Hastings 2020), relatively little ecological research has considered the scope of nature’s abiotic variability in maintaining species diversity (Abrams 2022).

Researchers have clearly argued that temporal variation (e.g., stochasticity) can promote species coexistence via fluctuation-dependent coexistence mechanisms (e.g., storage effect, relative nonlinearity) (Chesson 2000, 2018; Adler et al. 2006; Meyer et al. 2022). With these mechanisms, temporal niche differentiation enables coexistence between species with different competitive advantages (Angert et al. 2009; Mathias and Chesson 2013; Miller and Klausmeier 2017). Despite the broad focus on environmental variation, one specific class of variability that has been less-well explored, yet ought to provide a generalizable and analytically tractable entryway into coexistence theory for variable environments, is that of periodic fluctuations – that is, repeated and predictable fluctuations in environmental conditions. For examples, (Litchman and Klausmeier 2001) use fast/slow approximations to elegantly show that night/day oscillations in light can mediate competitive outcomes in phytoplankton – analytical solation that are tricky to garner from environmentally stochastic models.

Studies are beginning to suggest that periodic environments may have significant implications for species coexistence (Mathias and Chesson 2013; Miller and Klausmeier 2017; White and Hastings 2020). For example, temporally changing resource conditions, which fluctuate between seasons of high productivity to seasons of very little to no productivity (Fretwell 1972; Chesson and Huntly 1997), may favour different species at different times of a period (Armstrong and McGehee 1980; Litchman and Klausmeier 2001; Hastings 2012; Huntly et al. 2021). Similarly, it has been suggested that certain life history trade-offs and temporal differentiation in competing species’ performance may alter coexistence outcomes in the face of periodic environments (Litchman and Klausmeier 2001; McMeans et al. 2020). Recently, Mougi (2020) extended competition results from single periodicities to polyrhythms (i.e., multiple interacting periodicities) to show that the coupling of differently timed resource fluctuations may broaden the range of coexistence between diverse species that rely on limited resources. While these recent papers highlight the importance of periodic variation, the demand for a more general theoretical understanding on the role of periodic conditions – either in isolation or as suites of periodicities (i.e., polyrhythms) – remains (White and Hastings 2020; Abrams 2022). Notably, we lack a general understanding of coexistence in periodic environments akin to our well-established theoretical foundation framed around steady-state dynamics.

The development of such general theory is critical as climate change is currently altering the nature of these environmental fluctuations (Dijkstra et al 2011; Shuter et al. 2012; Urban et al. 2012; Chesson 2018; Al-Habahbeh et al. 2020). Northern-hemisphere winters are becoming shorter in length and more moderate (Caldwell et al. 2020; Edlund et al. 2017; Ficker et al. 2017; Warne et al. 2020), and weather patterns across the globe are becoming more variable and unpredictable (Fang and Stefan 1998; O’Reilly et al. 2015). In response, many communities have experienced an increase in species extinction (Urban et al. 2012; Moor 2017; Fung et al. 2020) and invasion rates (Stachowicz et al. 2002; Sharma et al. 2009; Dijkstra et al. 2011; Cerasoli et al. 2019; Atkinson et al. 2020). Therefore, as climate change continues to alter the abiotic conditions to which organisms have adapted to, the mechanisms regulating species coexistence may be fundamentally altered (di Paola et al. 2012; Korpela et al. 2013; Tunney et al. 2014; Anderson et al. 2015; Eloranta et al. 2016; Bartley et al. 2019; Caldwell et al. 2020). With all this in mind, developing an understanding for the mechanisms behind the maintenance of biodiversity in periodic environments becomes even more crucial.

Inspired by the fluctuating-light-driven coexistence results of Litchman and Klausmeier (2001), we sought to develop a generalizable framework for coexistence in fluctuating environments.

Towards this, we extend upon the seasonal coexistence model first introduced by McMeans et al. (2020) to more broadly explore the role of periodic forcing across time scales (e.g., days to multi-decadal) via different biological parameter combinations. We also generalized our approach by allowing high and low growth periods (not just high growth - no growth alone), with temporally-differential competitive abilities. Specifically, we extend the classical Lotka-Volterra coexistence criteria to include the role of environmental periodicities. This approach allows our results to be phrased around classical coexistence conditions with temporally-scaled inter- and intraspecific competition strengths. Here, we define a period as a unit of time that is composed of two distinct seasons of variable length and seek to generally explore which competitive outcomes may occur in these environments and under what biological conditions (i.e., different growth rates). Towards this, we employ an analytical approach consistent with the classical Lotka-Volterra phaseplane theory by developing a simple approximation that allows us to solve for the isocline solutions of a time-separated periodic model. Specifically, this approximation allows us to define coexistence criteria for periodic environments. We then illustrate how periodic environments can, in and of themselves, drive bifurcations (i.e., changing invasion criteria) such that competitive outcomes (i.e., stable coexistence, competitive exclusion, and contingent coexistence) are mediated by the environment. We end by discussing our competition results in light of how climate change is altering the nature of key underlying abiotic periodicities.

## Methods

We start by extending McMeans et al. (2020)’s annual seasonal model. Here, we define season more generally as a discrete division in time that repeats itself, or is periodic, of any given length within a period. As such, summer (more productive) and winter (less productive) seasons in McMeans et al. (2020) repeat themselves with a periodicity of one-year, but we may also similarly decompose other naturally shorter (e.g., seconds (Huntly et al. 2021)) and longer (e.g., El-Nino Southern Oscillations (Joseph and Kumar 2021)) periods of time into discrete seasons of more or less productive conditions. Towards this general understanding of periodic environments, we extend the Lotka-Volterra competition model (Chesson 2018) into a periodic model that repeatedly alternates between two discrete seasons, a productive (*f*_*p*_) and less productive (*f*_*LP*_) season. For each species, these functions are modelled with environmentally specific parameter combinations to incorporate biological constraints within each season, discussed below (Fig. 1a). The Lotka-Volterra model is defined as:

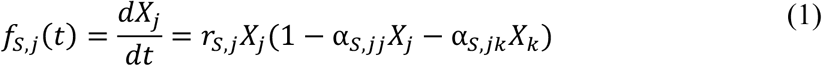

**Fig. 1.**
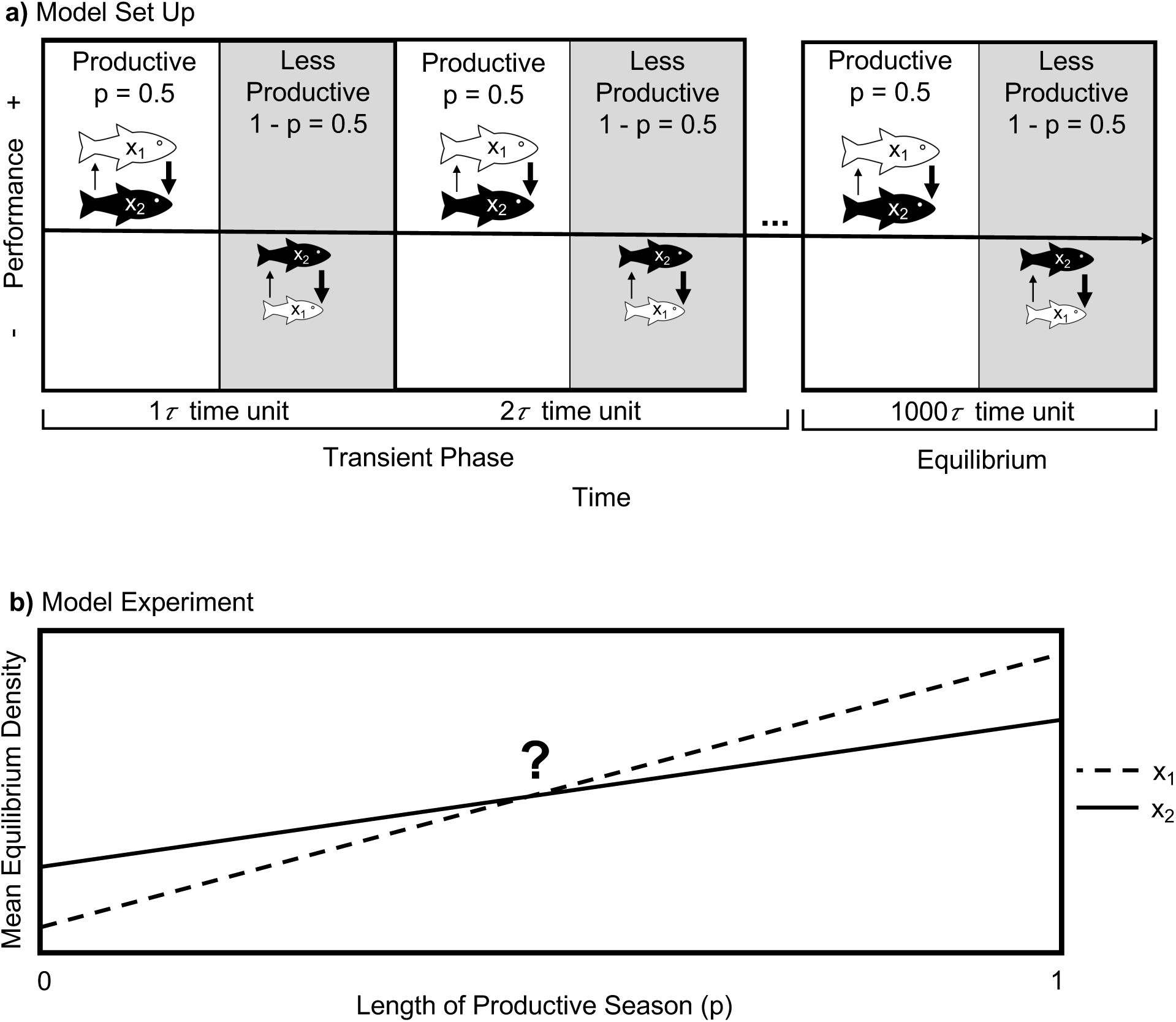
Model schematic investigates the projected mean equilibrium density over the length of the productive season when species compete under different periodic conditions. a) The model set-up investigates two time-separated seasons with different environmental conditions (productive (white box) and less productive (gray box)). In numerical simulations, competitive species, *X1* (white fish) and *X2* (black fish), exhibit different trade-offs in response to these differing environmental conditions, where each species may experience different levels of inter- to intraspecific competitive interactions (thickness of arrows) depending on which species may have an overall better performance (in terms of growth and competition) in one season compared to the other. An experiment is projected over many periods (say 1000 time-units) of length *τ* until it reaches an asymptotic state (we refer to this as an equilibrium despite the within-year variation). b) In the model experiment, the projected mean equilibrium density, which represents a fluctuating’ species density at its asymptote, is calculated as the duration of the productive season (as a proportion of each time unit) varies from 0 – 1.

**Fig. 2.**
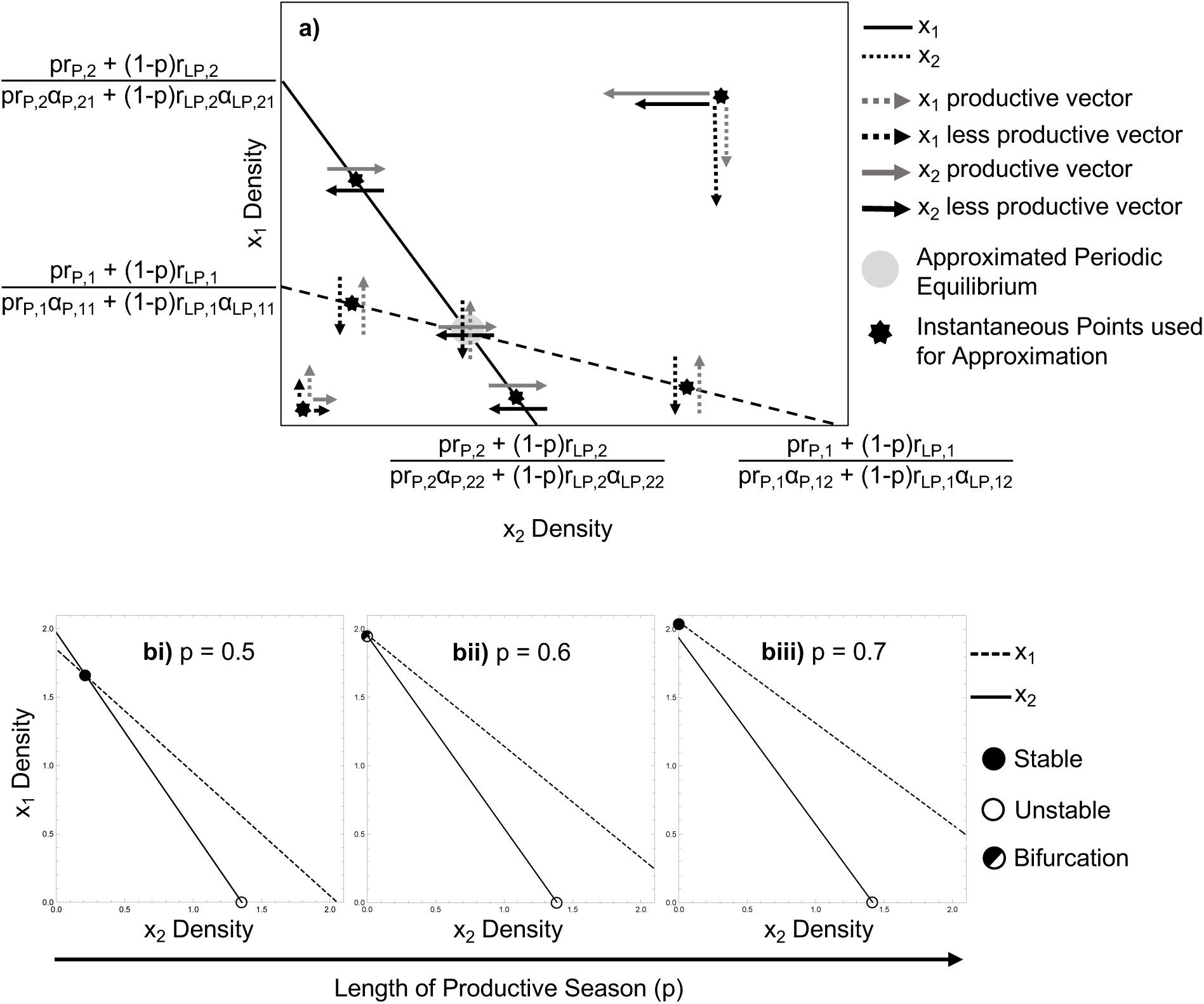
Isocline approximation of *X1* density on *X2* density. a) Stable coexistence isocline approximation on phaseplane of *X1* (dashed line) and *X2* (solid line) densities. Approximate isocline solutions for a given species occur when the instantaneous rate of change (represented by vector arrows) in each season, scaled by seasonal duration, are equal and opposite of each other (i.e., they “cancel” each other out such that the net change over a full period is null). Refer to Table 1 for similarities to Lotka–Volterra coexistence conditions. b i-iii) illustrates a *p*-driven bifurcation, showing the movement of the isoclines as the productivity season travels from i) *p* = 0.5; coexistence, to ii) *p* = 0.6; transcritical bifurcation, to iii) *p* = 0.7; competitive exclusion of species 2. Parametric values: *αP,11*=0.44, *αP,22*=0.66, *αP,12*=0.25, *αP,21*=0.53, *αLP,11*=1.11, *αLP,22*=0.83, *αLP,12*=1.83, *αLP,21*=0.48, *rP,1*=1.7, *rP,2*=1.2, *rLP,1*=0.3, *rLP,2*=1.

where *j* and *k* represent two competing species, and *S* represents a season (either productive, *P*, or less productive, *LP*). Here, *r*_*s,j*_ is the intrinsic rate of population growth for species *j* in a season, *S*, α_*s,jj*_ is the intraspecific competitive coefficient for species *j* in season *S*, and α_*S,jk*_ is the interspecific competitive coefficient describing the effect of species *k* on species *j* in season *S*.

When running simulations, and to maintain the spirit of the model assumptions (i.e., productive and less productive season), we assumed the following simple biologically realistic assumptions:

1. Maximal growth rates are larger in the productive season than the less productive season for both species (i.e., *r*_*P,j*_ > *r*_*LO,j*_), and;
2. Since resources are more available in the productive season compared to the less productive season, intraspecific competition will be lower in the productive season compared to the less productive season (i.e., α_*p,jj*_ < α_*LP,jj*_).

Further, to incorporate realistic biological trade-offs between competing species, we assumed that species 1 is a better performer (in terms of growth and competition) in the productive season compared to species 2, and the opposite is true in the less productive season. Keeping in mind the previous seasonal constraints, this produces the following realistic parametric trade-offs for the two species:

1. Species 1 has a higher growth rate in the productive season (i.e., *r*_*P,1*_) and a lower growth rate in the less productive season (i.e., *r*_*LP,2*_> *r*_*LP,*,1_) compared to species 2, and;
2. Species 1 has a smaller intraspecific competitive coefficient in the productive season (i.e., α_*LP,11*_ < α_*LP,22*_) and a larger intraspecific competitive coefficient in the less productive season (i.e., α_*LP,11*_ > α_*LP,22*_) compared to species 2.

These trade-offs are similar to an empirical case study by McMeans et al. (2020) where cold-adapted fish (e.g., lake trout, *Salvelinus namaycush*) are seasonal generalists with moderate growth rates year-round, and warm-adapted fish (e.g., smallmouth bass, *Micropterus dolomieu*), the lake trout’s competitor, are seasonal specialists with higher growth rates during the summer and lower growth rates during the winter. Note that we have not set any seasonal constraints on the interspecific competitive coefficients, therefore, α_#,%&_ can be any value.

With the above assumptions taken into consideration, our model is a periodic step function (repeats every 1-time unit, or period) that goes through a productive season (*P*), and a less productive season (*LP*), as follows:

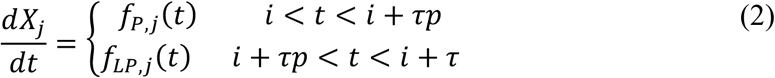

Where *i* is the period number (i.e. year) that runs on an integer step size of 1 from 0 to *t*_*prd*_(the number of periods that the model runs for). *τ* governs the length of each period, and *p* is the proportion of each period that is considered productive (i.e., τ*p* is the length of the productive season; leaving τ(*1-p*) as the length of the less productive season). The period length, defined by *τ*, allows us to examine how environmental periodicities, and associated biological trade-offs, across time scales may influence coexistence. As above, *j* represents one of the two competitive species, either species 1 or 2. These periodic functions of *f*_*p*_ and *f*_*LP*_ are defined in equation (1) as *f*_#_.

We coded all numerical simulations in Mathematica 12.0. The models are integrated over numerous periods until an asymptotic state, referred to hereafter as an equilibrium state, despite within-period variation, has been reached (i.e., mean value from 900 – 1000 time-units) to remove transient influences. However, within each time unit, as discussed above, they sequentially follow first productive then less productive parameters corresponding to the given productive seasonal fraction, *p*, within the period. The productive seasonal fraction, *p*, allows us to change the proportion of the period that is under our productive conditions versus our less productive conditions, (*1*-*p*) (e.g., increase *f*_!_) (Fig. 1b).

Finally, all model parameterizations for our simulations can be found in our figure legends and in the Supplementary Material. Below we first walk through our approximation and then our general analytical results before highlighting the generality of our analytical solutions using numerical simulations. For each outcome, we explore how changing the season lengths, via *p*, influences competitive coexistence and exclusion (Fig. 1b; Table 1).

**Table 1:** Lotka-Volterra Coexistence Conditions vs. Seasonal Coexistence Conditions.

|  | Lotka – Volterra | Seasonal Coexistence |
| --- | --- | --- |
|  | Criteria | Criteria |
| Stable Coexistence | $\alpha_{11} > \alpha_{21}$ | $\frac{\mathbf{p}r_{P,1}\alpha_{P,11} + (1 - \mathbf{p})r_{LP,1}\alpha_{LP,11}}{\mathbf{p}r_{P,1} + (1 - \mathbf{p})r_{LP,1}} > \frac{\mathbf{p}r_{P,2}\alpha_{P,21} + (1 - \mathbf{p})r_{LP,2}\alpha_{LP,21}}{\mathbf{p}r_{P,2} + (1 - \mathbf{p})r_{LP,2}}$ |
| | $\alpha_{22} > \alpha_{12}$ | $\frac{\mathbf{p}r_{P,2}\alpha_{P,22} + (1 - \mathbf{p})r_{LP,2}\alpha_{LP,22}}{\mathbf{p}r_{P,2} + (1 - \mathbf{p})r_{LP,2}} > \frac{\mathbf{p}r_{P,1}\alpha_{P,12} + (1 - \mathbf{p})r_{LP,1}\alpha_{LP,12}}{\mathbf{p}r_{P,1} + (1 - \mathbf{p})r_{LP,1}}$ |
| Competitive Exclusion by $sp_1$ | $\alpha_{11} > \alpha_{21}$ | $\frac{\mathbf{p}r_{P,1}\alpha_{P,11} + (1 - \mathbf{p})r_{LP,1}\alpha_{LP,11}}{\mathbf{p}r_{P,1} + (1 - \mathbf{p})r_{LP,1}} > \frac{\mathbf{p}r_{P,2}\alpha_{P,21} + (1 - \mathbf{p})r_{LP,2}\alpha_{LP,21}}{\mathbf{p}r_{P,2} + (1 - \mathbf{p})r_{LP,2}}$ |
| | $\alpha_{22} < \alpha_{12}$ | $\frac{\mathbf{p}r_{P,2}\alpha_{P,22} + (1 - \mathbf{p})r_{LP,2}\alpha_{LP,22}}{\mathbf{p}r_{P,2} + (1 - \mathbf{p})r_{LP,2}} < \frac{\mathbf{p}r_{P,1}\alpha_{P,12} + (1 - \mathbf{p})r_{LP,1}\alpha_{LP,12}}{\mathbf{p}r_{P,1} + (1 - \mathbf{p})r_{LP,1}}$ |
| Competitive Exclusion by $sp_2$ | $\alpha_{11} > \alpha_{21}$ | $\frac{\mathbf{p}r_{P,1}\alpha_{P,11} + (1 - \mathbf{p})r_{LP,1}\alpha_{LP,11}}{\mathbf{p}r_{P,1} + (1 - \mathbf{p})r_{LP,1}} > \frac{\mathbf{p}r_{P,2}\alpha_{P,21} + (1 - \mathbf{p})r_{LP,2}\alpha_{LP,21}}{\mathbf{p}r_{P,2} + (1 - \mathbf{p})r_{LP,2}}$ |
| | $\alpha_{22} < \alpha_{12}$ | $\frac{\mathbf{p}r_{P,2}\alpha_{P,22} + (1 - \mathbf{p})r_{LP,2}\alpha_{LP,22}}{\mathbf{p}r_{P,2} + (1 - \mathbf{p})r_{LP,2}} < \frac{\mathbf{p}r_{P,1}\alpha_{P,12} + (1 - \mathbf{p})r_{LP,1}\alpha_{LP,12}}{\mathbf{p}r_{P,1} + (1 - \mathbf{p})r_{LP,1}}$ |
| Contingent Coexistence | $\alpha_{11} < \alpha_{21}$ | $\frac{\mathbf{p}r_{P,1}\alpha_{P,11} + (1 - \mathbf{p})r_{LP,1}\alpha_{LP,11}}{\mathbf{p}r_{P,1} + (1 - \mathbf{p})r_{LP,1}} < \frac{\mathbf{p}r_{P,2}\alpha_{P,21} + (1 - \mathbf{p})r_{LP,2}\alpha_{LP,21}}{\mathbf{p}r_{P,2} + (1 - \mathbf{p})r_{LP,2}}$ |
| | $\alpha_{22} < \alpha_{12}$ | $\frac{\mathbf{p}r_{P,2}\alpha_{P,22} + (1 - \mathbf{p})r_{LP,2}\alpha_{LP,22}}{\mathbf{p}r_{P,2} + (1 - \mathbf{p})r_{LP,2}} < \frac{\mathbf{p}r_{P,1}\alpha_{P,12} + (1 - \mathbf{p})r_{LP,1}\alpha_{LP,12}}{\mathbf{p}r_{P,1} + (1 - \mathbf{p})r_{LP,1}}$ |

## Results

Here, we present our approximated isocline solutions and investigate the resulting coexistence criteria and behaviour under seasonality. Next, we reveal that changing season length, *p*, in response to climate change, mediates coexistence, competitive exclusion, and contingent coexistence (Fig. 3). Finally, we explore the robustness of our seasonally-mediated outcomes (i.e., seasonally-driven bifurcations) across a range of period lengths (Figs. 4-6).

**Fig. 3.**
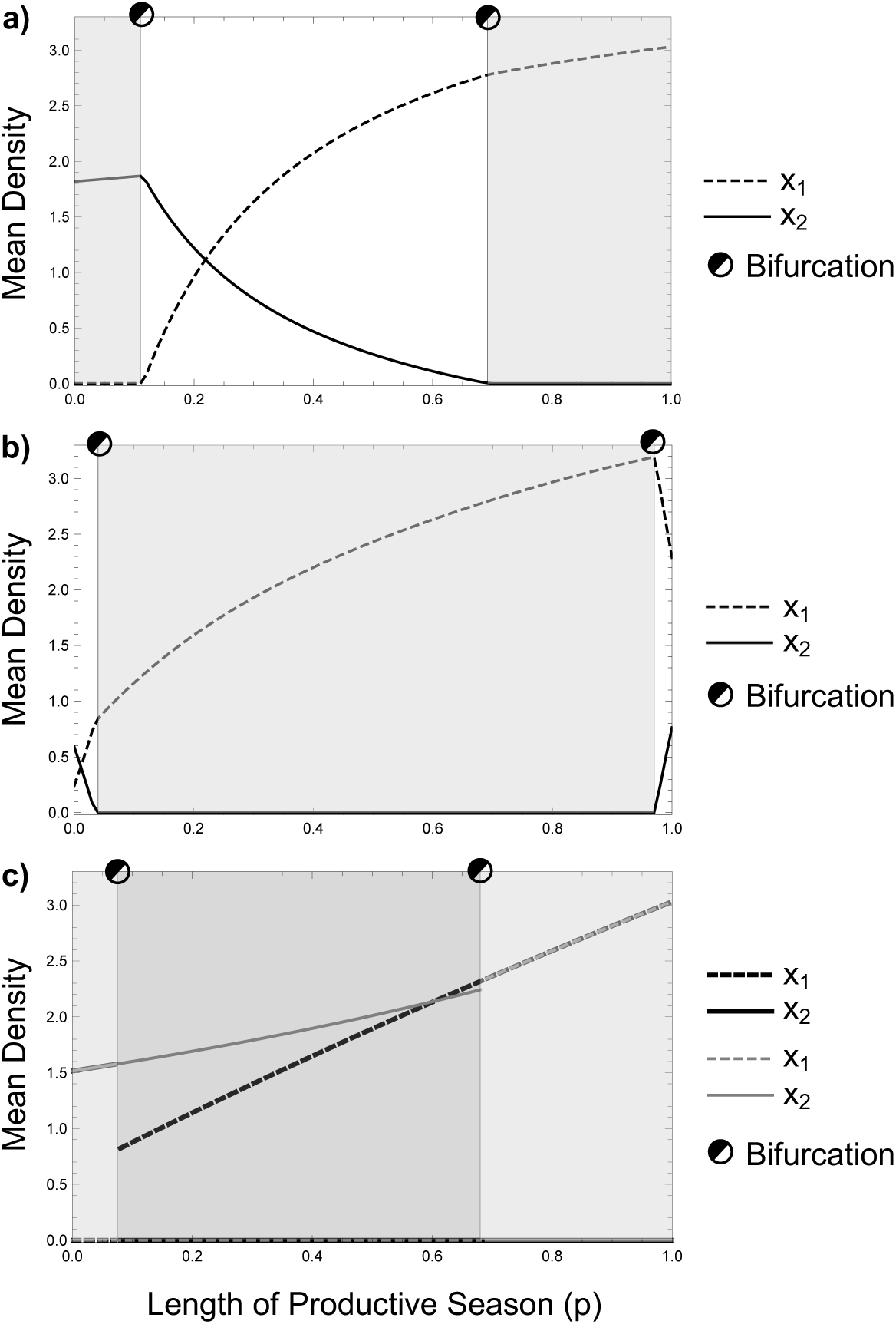
Numerical simulations of mean density over length of productive season (*p*) for seasonally-mediated outcomes. White zone represents stable coexistence, light gray zone represents competitive exclusion, and dark gray zone represents contingent coexistence where the equilibrium is unstable and multiple attractors exist between the two species. a) Seasonally- mediated coexistence. Parametric values: *αP,21*=0.35, *αP,12*=0.165, *αP,11*=0.33, *αP,22*=0.436, *rP,1*=1.7, *rP,2*=1.2, *αLP,21*=0.385, *αLP,12*=0,805, *αLP,11*=0.73, *αLP,22*=0.55, *rLP,1*=0.3, *rLP,2*=1. b) Seasonally-mediated competitive exclusion. Parametric values: *αP,21*=0.29, *αP,12*=0.38, *αP,11*=0.31, *αP,22*=0.44, *rP,1*=4, *rP,2*=1.2, *αLP,21*=1.27, *αLP,12*=1.02, *αLP,11*=1,67, *αLP,22*=1.18 *rLP,1*=0.3, *rLP,2*=1. c) Seasonally-mediated contingent coexistence. Parametric values; *αP,21*=0.548, *αP,12*=0.33, *αP,11*=0.33, *αP,22*=0.363, *rP,1*=1.7, *rP,2*=1.2, *αLP,21*=1.31, *αLP,12*=1.89, *αLP,11*=1.65, *αLP,22*=0.66, *rLP,1*=0.3, *rLP,2*=1.

**Fig. 4.**
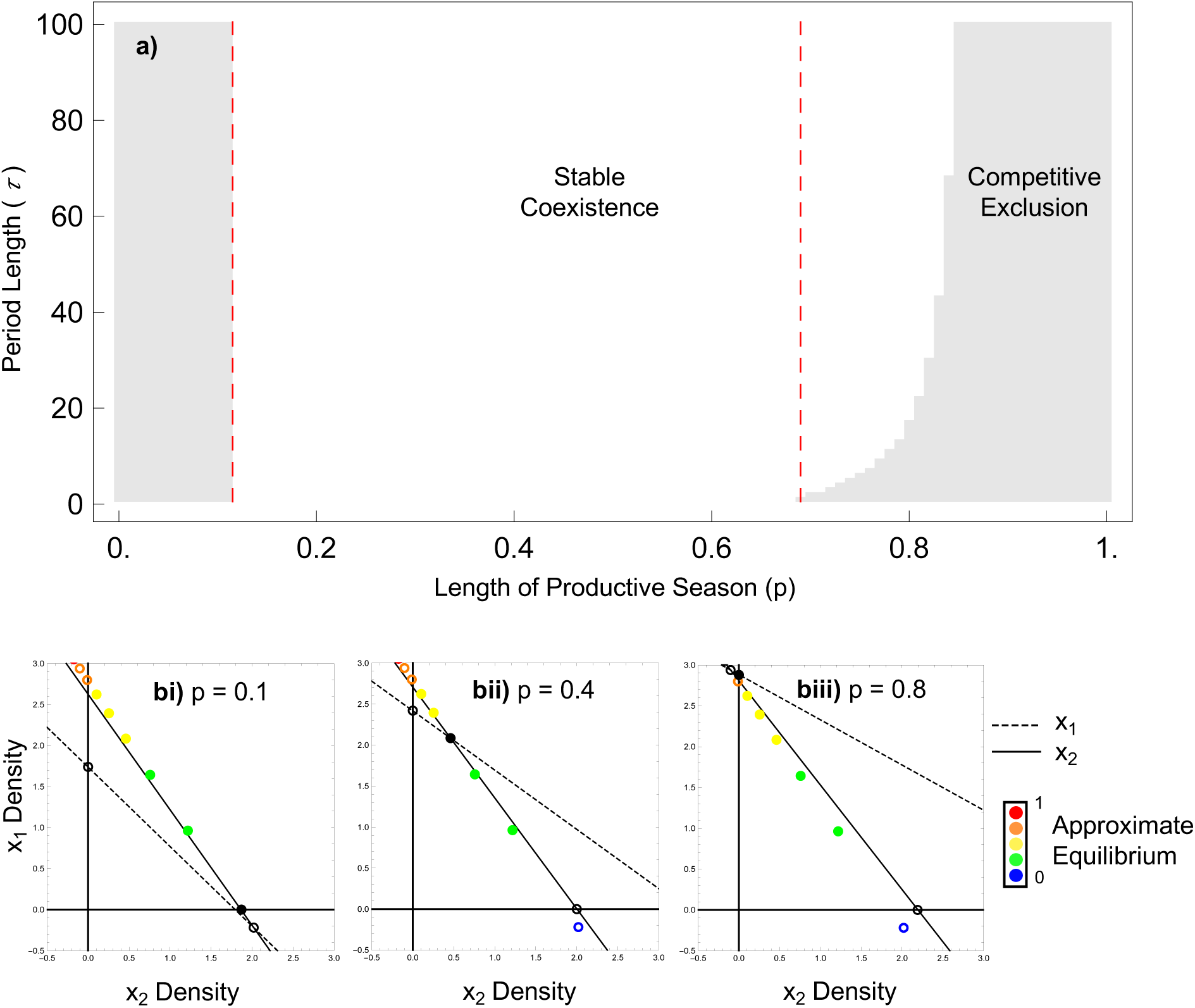
Two-dimensional bifurcation diagram of seasonally-mediated coexistence. a) seasonally- mediated coexistence expanding over a wide range of period lengths (*τ)*. The red dashed lines represent the isocline approximation’s prediction of the two transcritical bifurcation points at *p*=0.115 and *p*=0.69 (see Fig. S3.1 for complete transition across *p*, and S4.1 for explanation of approximation accuracy of seasonally mediated coexistence). b) isocline approximation tracks the approximate equilibrium, which represents the mean asymptotic behavior, as the productive season (*p*) increases from i) *p*=0.1; competitive exclusion of species 1, to ii) *p*=0.4; stable coexistence, to iii) *p*=0.8; competitive exclusion of species 2. Filled in circles are stable equilibrium points, and open circles are unstable equilibrium points. Parametric values: *αP,21*=0.35, *αP,12*=0.165, *αP,11*=0.33, *αP,22*=0.436, *rP,1*=1.7, *rP,2*=1.2, *αLP,21*=0.385, *αLP,12*=0,805, *αLP,11*=0.73, *αLP,22*=0.55, *rLP,1*=0.3, *rLP,2*=1.

### Approximate Isocline Solutions for the Periodic Lotka-Volterra Model

Although we use a periodically forced system, we can use equilibrium concepts to understand the dynamics of our model. Specifically, our model reaches an attractor such that the densities fluctuate modestly up and down on the attractor around a mean that does not change (i.e., an asymptotic state or dynamic equilibrium; Fig.S2.2). Given this equilibrium-like dynamic, we are interested in considering this asymptotic behaviour in a manner similar to the way we would for a system that reaches a true equilibrium. Here, we provide an approximation for our model isoclines and equilibria that mirror those of the original Lotka-Volterra competition model.

Note that the isoclines can be solved by recognizing that each period of *τ* time units would necessarily have to result in zero net growth (i.e., no overall changes in density between the start and end of each period), akin to classical zero-growth isoclines with a deterministic equilibrium.

That is, from System (2), the *Xj* zero-net-growth condition occurs when 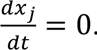. From this, the isocline solution follows as:

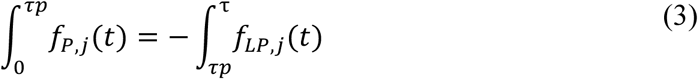

This isocline solution (3) is not analytically tractable (note that each season has dynamics over a time interval), making it seem as though an isocline approach appears infeasible.

Towards solving for an approximation for periodically forced isoclines, we take a slightly different approach then the classical linearization of the full equations (see Litchman and Klausmeier (2001) for an example). As our model is not tractable with this approach, we instead proceed by assuming that we can find a linearized approximation to the isocline from the periodic model by searching for points in phase space where the linearization of each seasons’ dynamics are exactly negated by each other (for both the *X1* and *X2* isoclines). We define such points as **zero net growth** points in the phaseplane (i.e., a point on the forced model’s isocline) and we define the collection of these points as **the linearized isocline approximation of the periodically-forced model**. Clearly, as the length of the period, *τ*, goes to 0, the error in this linear approximation will also go to 0. However, for longer periods or large growth rates (*r*), non-linear dynamics may drive this simplification to work poorly (see Supplementary Material

S4). Because of this, we numerically check our analytical results with numerical calculations throughout. We note that despite this, the results work surprisingly well across large parameter values suggesting the linear approximation works even when some degree of nonlinear dynamics are expressed.

By assuming that the instantaneous rates of change for each species at this *X1-X2* co-ordinate are linear, we approximate the dynamics over the time period, *τ*, by solving the equations using the Fundamental Theorem of Calculus over each period (see Supplementary Material S1). That is, over the interval fraction, *τp*, the productive trajectory scales linearly on the co-ordinate *X1-X2* as:

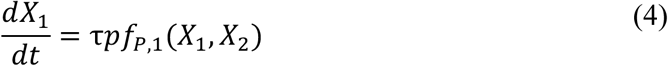

and the less productive trajectory over the interval fraction *τ*(*1-p*) scales linearly as:

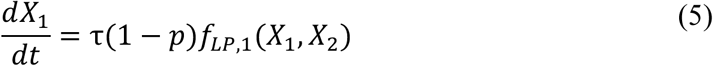

Given Equations (4) and (5), then the linearized dynamics at a *X1-X2* co-ordinate negate each other (i.e., resulting in the *X1*-isocline) when:

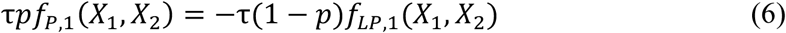

This approximation (Equation (6)), if it works, can be solved symbolically as done elegantly for the classical Lotka-Volterra model and thus allows an entry point into well-known coexistence analyses and interpretations.

From Equation (6), we see that *τ* factors out. After substituting the productive and less productive models of Equation (1) into Equation (6) and then some algebra, the resulting isocline solutions, for both species, follow the following form:

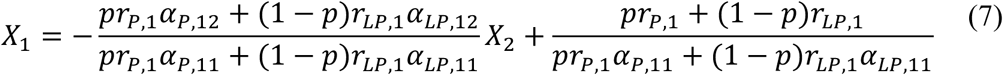

The geometry of the approximated isoclines (Equation (7)) resembles that of the original Lotka-Volterra isoclines, which therefore allows us to identify general rules for coexistence in periodic environments akin to the original coexistence criteria (Fig. 2a). Here, the isocline approximation solutions (and therefore coexistence criteria) are now based on temporal-growth-scaled inter- versus intra-specific competition rates (whereas the original Lotka-Volterra conditions are not dependent on growth rates; see Table 1 for Lotka-Volterra vs. Seasonal Coexistence Criteria).

Thus, the approximation enables us to generally determine and graphically explore how periodic environments and biological rates may interact in regulating competitive outcomes (Fig. 2b).

Immediately, based on these conditions, we can see that the duration of each season, and the corresponding biological conditions within them, play a critical role in regulating competitive outcomes in periodic environments. That is, our parameter *p* may drive bifurcations, and therefore alter competitive outcomes under changing climatic conditions (Fig. 2b).

We found numerically that this approximation works quite well (see Supplementary Material S2 for examples) and was, for example, able to repeatedly and accurately calculate the transcritical bifurcation points where an interior equilibrium intersects and exchanges stability, with an axial solution. Other approaches similar to our approximation, such as Litchman and Klausmeier (2001), have been used to accurately investigate species coexistence when organismal dynamics are fast relative to their forcing speed. Given our assumption of linearity, the relative time scale of forcing vs. dynamics ought to be important for our approximation’s accuracy. Indeed, when either the period length (*τ*) or the growth rate (*r*) becomes larger, the apparent length of each period increases, allowing for more nonlinearity within each season’s dynamics, which results in the approximation to fail at predicting the asymptotic behaviour (see Supplementary Material S4). Nevertheless, the range at which this approximation holds is impressively robust.

### General outcomes of changing environments

Given that changing season lengths may have powerful implications for species coexistence, we now ask how changing season length, *p*, (i.e., altering seasonal asymmetry) affects competitive outcomes. To do this we vary the season length from *p*=0 (i.e., the full period being entirely less productive) to *p*=1 (i.e., the full period being entirely productive). We note that these endpoints (i.e., *p*=0 or *p*=1) are constant (i.e., no seasonality) and thus, they reduce to the classic Lotka-Volterra coexistence criteria based on the individual biological conditions for *P* or *LP* (Table 1). These endpoints also give us reference conditions for exploring the effects of environmental periodicity and seasonally-mediated coexistence outcomes.

Since McMeans et al. (2020) found the intriguing case of seasonally-mediated coexistence, we were interested in using our analytical results to unpack other examples of how seasonal change could fundamentally alter competitive outcomes. McMeans et al. (2020) noted that where the boundary conditions yielded competitive exclusion (i.e., exclusion at *p*=0 and *p*=1), a seasonal model could produce coexistence for intermediate *p*-values. This is intriguing as it immediately suggests that alterations in *p* (say from climate change) can drive exclusion. Importantly, our analytical solutions (Table 1) suggest that seasonality can mediate all possible competitive outcomes at intermediate *p*-values (Fig. 3). Indeed, we found seasonally-mediated coexistence, competitive exclusion, and contingent coexistence (i.e., alternative states) (Fig. 3a-c respectively).

For these seasonally-mediated outcomes, when the qualitative competitive outcome at intermediate *p* is fundamentally different from the extremes (i.e., *p* = 0 and 1), *p* drives a series of transcritical bifurcations that move the system between different conditions in Table 1 (shifts between shaded and clear zones indicate transcritical bifurcations that alter the qualitative nature of the attractor in Fig. 3). Specifically, under our given sets of parametric combinations, changing *p* always drives a series of two bifurcations for these seasonally-mediated outcomes (Fig. 3). Other competitive outcomes of course are seasonally sensitive as they result in *p*-driven bifurcations when coexistence outcomes transition between the two endpoint conditions.

Additionally, we note that these transcritical bifurcations can be considered as changes in the invasion criteria – in this case, seasonally-altered invasion criterion – as they are commonly referred to in competition theory. For example, in Fig. 3a, the gray shaded regions indicate that one species cannot invade from small densities while in the white or unshaded region, it suddenly can invade from small densities.

Finally, clearly not all parameter combinations are sensitive to *p*-driven bifurcations. In these cases, *p* may simply drive changes in species’ densities rather than shifts in the equilibrium structure (e.g., as seen in McMeans et al. (2020)). While here we broadly unpack *p*-driven bifurcations to explore the generality of these outcomes, we note that all results below have nearby solutions that do not undergo bifurcations yet and are qualitatively similar in terms of the general effect of *p* on the isocline and equilibrium geometry.

### Robustness of Seasonally-Mediated Competitive Outcomes

To fully understand how robust the different seasonally-mediated outcomes are we looked at the bifurcation structure in 2-dimensional parameter space, by using both season length, *p*, (as in Fig. 3) and the period length, *τ*, as bifurcation parameters. We note that if seasonally-mediated outcomes exist across a broad range of period lengths, *τ*, then this suggests that seasonally- mediated competitive outcomes could be found across a wide range of natural periodicities (from diurnal to multi-decadal).

In Fig. 4a, we document cases where a lack of periodicity (i.e., at *p*=0 or 1) drives competitive exclusion while for a broad range of *τ*’s (ranging from 1-100 time units), we have species coexistence at intermediate *p*-values. Our analytical solutions allow us to track the “mean equilibrium” through each transcritical bifurcation from competitive exclusion of species 1 to species coexistence and finally to competitive exclusion of species 2 as the length of the productive period, *p*, increases (i.e., changing *p* shifts between competitive exclusion and stable coexistence conditions in Table 1; Figs. 4bi-iii, S3.1). The robustness of this pattern implies that for any given *τ*, seasonality alone can mediate coexistence at intermediate *p*-values. As the period length (*τ*) grows, the numerically-generated region of coexistence broadens, and the second transcritical bifurcation point, the transition between species coexistence and competitive exclusion of species 2, deviates away from the analytically-determined transcritical bifurcation point (see more regarding approximation accuracy of seasonally mediated coexistence in S4.1). This suggests that the analytical bifurcation structure is sensitive to different parametric values (which is also suggested by the seasonal coexistence criteria shown in Table 1), but nonetheless, the result of seasonally-mediated coexistence is qualitatively general. We note that the simple linear prediction breaks down as r*τ* grows (i.e., larger *τr* allows the nonlinear dynamics to express themselves; see Supplementary Material S4.1). Thus, we see that seasonally-mediated coexistence occurs very broadly suggesting that a large range of natural periodicities may be powerful drivers of coexistence.

Similarly, in Fig. 5a, we document cases where species coexistence occurs in static environments (i.e., at *p*=0 or 1), while competitive exclusion occurs at intermediate *p-*values over a broad range of *τ*’s (here, from 1-40 time units). Following McMeans et al. (2020)’s naming convention, we refer to this as seasonally-mediated competitive exclusion. As in the above case, with increasing *p*, our analytical approximation tracks the isoclines and the “mean equilibrium” from stable coexistence to competitive exclusion of species 2 through a transcritical bifurcation on the *X2*=0 axis, and finally back through the axis (via another bifurcation) to stable coexistence (Figs. 5b, S3.2). The transcritical bifurcations points determined by our analytical solution accurately track the numerical solutions over several period length values of *τ* before the seasonal bifurcations cease at higher values of *τ*. Again, we note that the simple linear prediction breaks down as *τr* grows (i.e., larger *τr* allows the nonlinear dynamics to express themselves; see Supplementary Material S4.2). The 2-dimensional bifurcation diagram again shows a broad range in *τ*-values that yield competitive exclusion; however, it does not stretch across all *τ*-values (i.e., very large period lengths). While the exact range of parameter space that this seasonally-mediated competitive exclusion occurs at is clearly dependent on other parameters, this result is still quite general over a wide range of *τ*-values, suggesting that periodic environments may also be powerful drivers of competitive exclusion. Here, as *τ* grows, this tongue of competitive exclusion slowly shrinks across intermediate values of *p*. Again, this is an interesting result, which suggests that simple alterations in season length may fundamentally alter competitive outcomes.

**Fig. 5.**
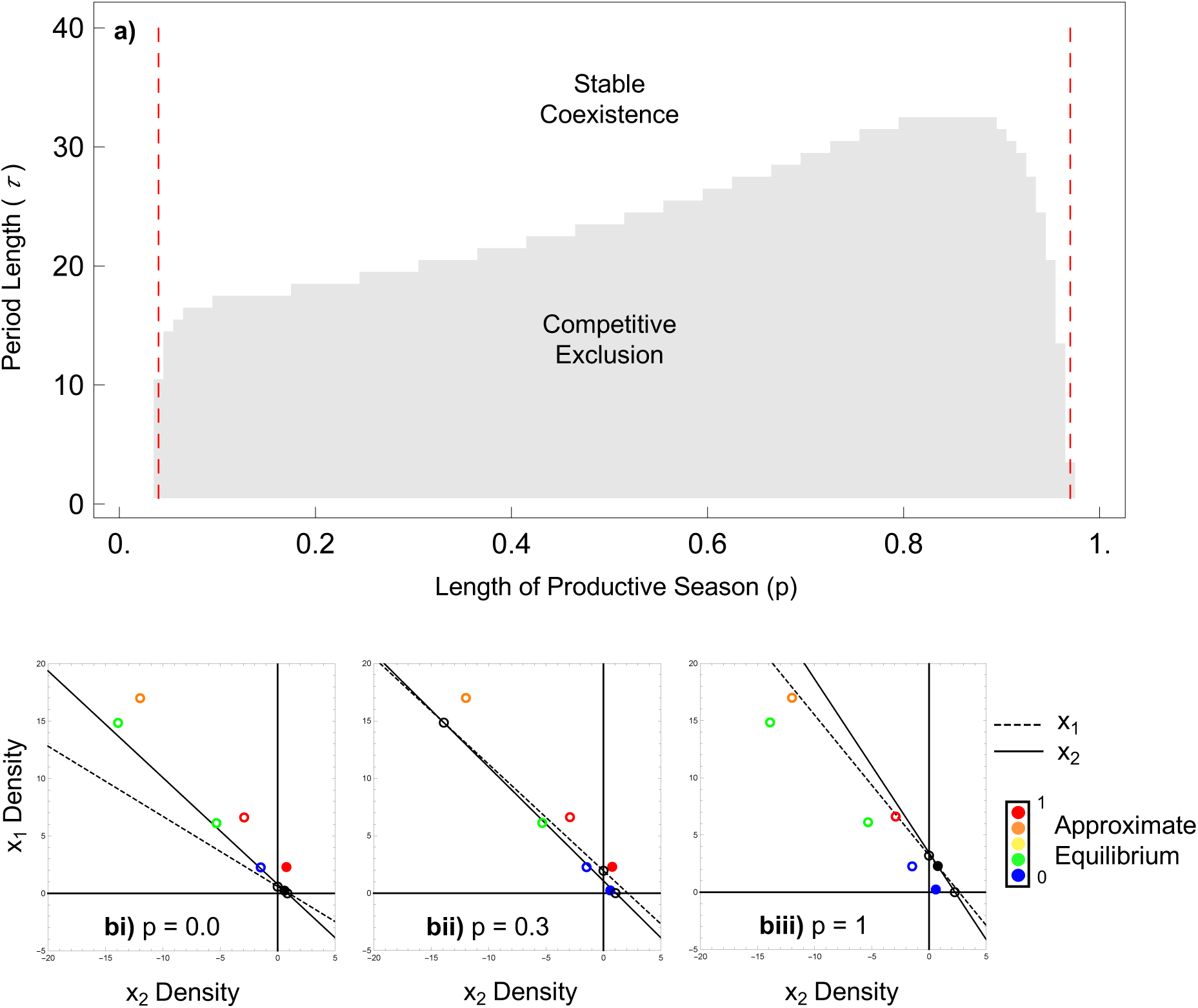
Two-dimensional bifurcation diagram of seasonally-mediated competitive exclusion. a) seasonally-mediated competitive exclusion expanding over a wide range of period lengths (*τ*). The red dashed lines represent the isocline approximation’s prediction of the two transcritical bifurcation points at *p*=0.04 and *p*=0.97 (see Fig. S3.2 for complete transition across *p*, and S4.2 for explanation of approximation accuracy of seasonally mediated competitive exclusion). b) isocline approximation tracks the approximate equilibrium, which represents the mean asymptotic behaviour, as the productive season increases from i) *p*=0.0; stable coexistence, to ii) *p*=0.3; competitive exclusion of species 2, and back to iii) *p*=1.0; stable coexistence. Filled in circles are stable equilibrium points, and open circles are unstable equilibrium points. Parametric values: *αP,21*=0.29, *αP,12*=0.38, *αP,11*=0.31, *αP,22*=0.44, *rP,1*=4, *rP,2*=1.2, *αLP,21*=1.27, *αLP,12*=1.02, *αLP,11*=1,67, *αLP,22*=1.18 *rLP,1*=0.3, *rLP,2*=1.

Finally, in Fig. 6a, similar to the above seasonally-mediated coexistence case with competitive exclusion at the boundaries (i.e., *p*=0 or 1), we now document cases where contingent coexistence (i.e., alternative states) occurs at intermediate *p* values across a broad range of *τ*’s (here, from 1-100 time units). Although this outcome is quite similar to seasonally-mediated coexistence with competitive exclusion being located at each boundary (i.e., *p*=0 or 1), its isocline geometry, based off of parametric combinations, results in seasonally-mediated contingent coexistence at intermediate *p*-values (Figs. 6bi-iii, S3.3). Our approximation tracks the analytical non-trivial (interior) equilibrium, which in this case is unstable in positive (*X1, X2*) state space (Fig. 6bii). Here, initial densities determine which species will dominate and which species will become extinct with time (i.e., which of the two stable axial solutions, shown in Fig. 6bii, the system will end up at). The robustness of this pattern also implies that for any given *τ*, seasonality may be a powerful driver of contingent coexistence (i.e., changing *p* moves from competitive exclusion to contingent coexistence conditions in Table 1). As *τ* grows, the numerically-generated contingent coexistence region expands across higher productive season lengths before it narrows around lower lengths of the productive season. Again, our analytical solution accurately determines at which *p* values the transcritical bifurcations will occur for very small *τ* values, but falls off with larger period lengths, *τ* (see Supplementary Material S4.3 for approximation accuracy of seasonally mediated contingent coexistence), despite the general phenomenon remaining across time scales.

**Fig. 6.**
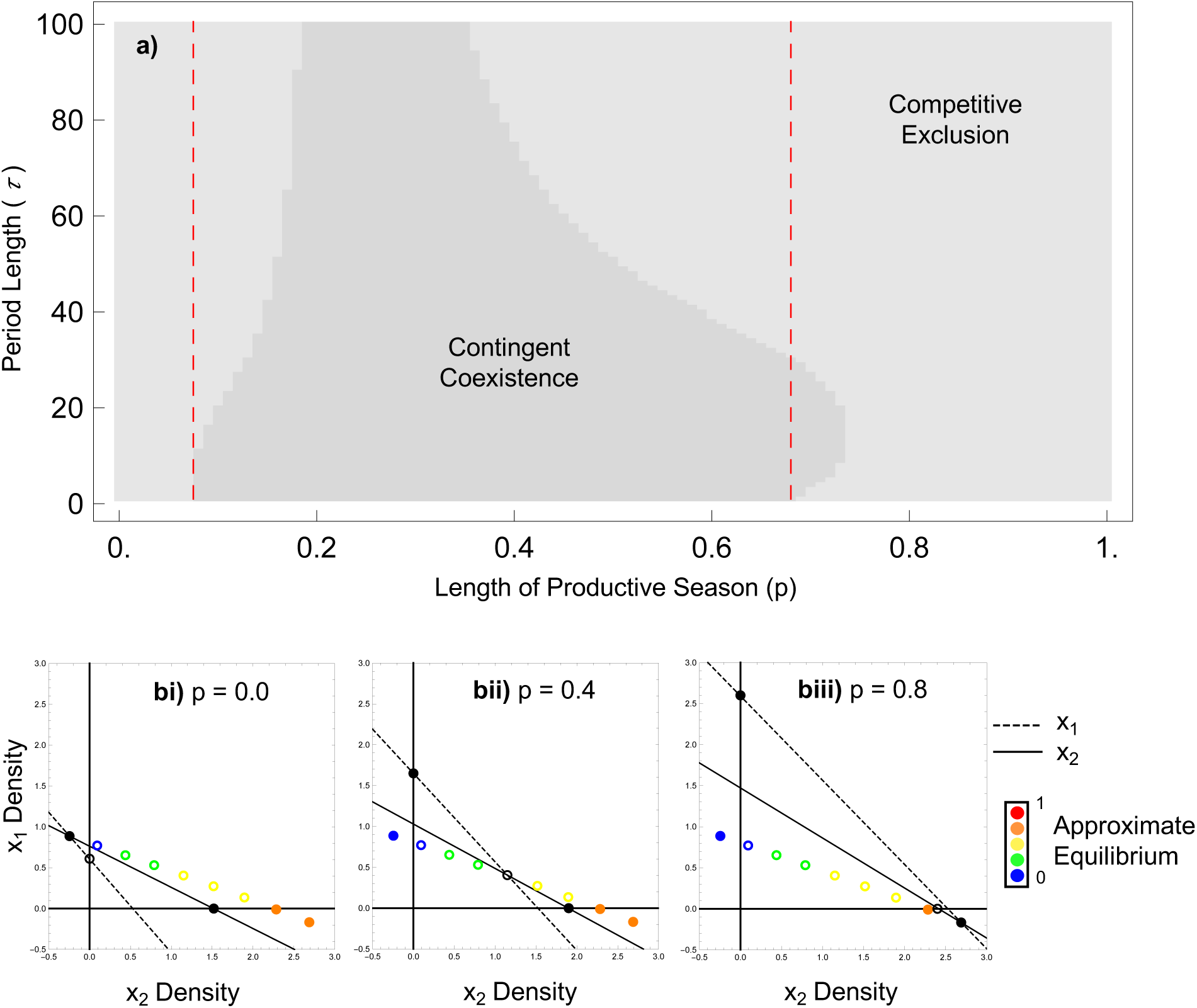
Two-dimensional bifurcation diagram of seasonally-mediated contingent coexistence. a) seasonally-mediated contingent coexistence expanding over a wide range of period length (*τ*). The red dashed lines represent the isocline approximation’s prediction of the two transcritical bifurcation points at *p*=0.07 and *p*=0.69 (see Fig. S3.3 for complete transition across *p*, and S4.3 for explanation of approximation accuracy of seasonally mediated contingent coexistence). b) isocline approximation tracks the approximate equilibrium, which represents the mean asymptotic behaviour, as the productive season increases from i) *p*=0.0; competitive exclusion of species 1, to ii) *p*=0.4; contingent coexistence, to iii) *p*=0.8; competitive exclusion of species 2. Filled in circles are stable equilibrium points, and open circles are unstable equilibrium points. Parametric values; *αP,21*=0.548, *αP,12*=0.33, *αP,11*=0.33, *αP,22*=0.363, *rP,1*=1.7, *rP,2*=1.2, *αLP,21*=1.31, *αLP,12*=1.89, *αLP,11*=1.65, *αLP,22*=0.66, *rLP,1*=0.3, *rLP,2*=1.

These three seasonally-mediated outcomes suggest that seasonal alterations, due to climate change, may drive precipitous changes in competitive systems. All of these seasonally-mediated outcomes appear general and may be found across various ecosystems that experience abiotic fluctuations of any period length. Similarly, for a given period length (e.g., annual variation), these outcomes may be highly general for competing organisms with a range of life history strategies (i.e., vital rates that dictate the speed of biotic dynamics relative to any environmental variation). On the other hand, depending on parametric values, not all instances of periodic variation will drive these series of bifurcations. Under some conditions, there are simply *p*-driven changes in species densities between the boundary conditions where only one bifurcation (Figs. S2.1e, S2.1f) or no bifurcations (Figs. S2.1d) are found under changing periodicities.

## Discussion

Here, we present a linear approximation similar in approach to other periodic models (e.g., Han et al. (1999); Litchman and Klausmeier (2001)) to analytically solve for a periodic Lotka-Volterra competition model (see Table 1 for seasonal coexistence criteria). Our linear approximation quantitatively breaks down for large values in *τr* but still largely operates to qualitatively predict the nature of periodic-forcing on competition (e.g., Fig. 4a,6a). This approximation, akin to the longstanding classical Lotka-Volterra coexistence conditions, importantly allows us to derive a parallel set of coexistence conditions for periodic environments. With the inclusion of periodic environmental forcing, we find that species coexistence depends on seasonal-growth-scaled inter- versus intra-specific competition strengths (Table 1). That is, where classical stable coexistence requires intraspecific competition to be greater than interspecific competition, for both species, we show that intraspecific competition, scaled by seasonal growth, must be greater than interspecific competition, scaled by seasonal growth, for both species.

These seasonal coexistence conditions importantly suggest that species’ coexistence may be incredibly sensitive to changes in season length. While McMeans et al. (2020) found seasonally- mediated coexistence through empirically-motivated numerical simulations, our results show that seasonality can mediate coexistence, competitive exclusion, and contingent coexistence (Fig. 3). That is, seasonality, in and of itself, can drive all possible competitive outcomes (i.e., Table 1).

As these seasonally-mediated outcomes appear across a vast range of period lengths (Fig. 4-6), this suggests that these seasonally-mediated outcomes are quite robust, and can be found across a wide range of different temporal scales and life history strategies (i.e., fast vs. slow life history strategies).

As the change in season length (*p*) may have an important role in mediating competitive outcomes (Fig. 3), climate change is altering environmental periodicities across a range of temporal scales (Dijkstra et al. 2011; Shuter et al. 2012; Urban et al. 2012; Chesson 2018; Al- Habahbeh et al. 2020), and our results suggest that this could have sudden and drastic – sometimes unpredictable – effects on coexistence. Often the responses to changing seasons are nonlinear, and some of these outcomes are unexpected based on the classical Lotka-Volterra conditions under non-seasonal environments (e.g., nonlinear paths between the boundary conditions at *p* = 0 and 1; Figs. 3b, 5, S2.1b and d). As an example, we find cases where competitive exclusion occurs under intermediate season lengths, even though coexistence is expected under the same non-seasonal conditions (i.e., coexistence occurs for entirely low- or entirely high-productive conditions; Fig. 3b). Alarmingly, these strong nonlinear effects of changing season length, *p*, suggest that precipitous changes in species density and composition may unexpectedly occur as climate change alters the nature of seasonality.

Although simple, the seasonal competition model we examined here is based on biologically realistic assumptions about competitive species in temporally varying environments. First, periodic environments are commonly reflected by times of high and low productivity, where species tend to flourish during the productive seasons, and decline (e.g., mortality-dominated) during the less productive seasons as resource availability becomes sparse (Fretwell 1972;

Litchman and Klausmeier 2001; Mutze 2009; Klausmeier 2010; Hastings 2012; Lyu et al. 2016; Vihtakari et al. 2016). These periods therefore impact species’ growth rates, but also offer the potential for differential responses of competing species to seasonally-driven environmental variation (Chesson and Huntly 1997; Forrest and Miller-Rushing 2010; Shuter et al. 2012; Gao et al. 2016; Chesson 2018; Huang et al. 2019). It is well known that competing species can display temporal trade-offs in stochastic environments that can promote coexistence (Angert et al. 2009; Shuter et al. 2012; Mougi 2020). For instance, Litchman and Klausmeier (2001) discovered that slow fluctuations in light promote stable coexistence between competing species of phytoplankton who exhibit trade-offs in performance across temporally changing resource availability. More recently, and consistent with our theoretical results here, empirical evidence is beginning to suggest species may show trade-offs to regular seasonal fluctuations that might promote coexistence. For example, a species may be a seasonal specialist (e.g., display a very high growth rate during the summer and a low growth rate during the winter) while its competitor may be more of a temporal generalist, who’s able to maintain roughly the same growth rate year-round (Niiyama 1990; Chan et al. 2009; Abrams et al. 2013; Korpela et al. 2013; Meyer et al. 2022).

While we concentrate on temporal forcing in our model, seasonal signals in competition may often be related to spatial-temporal and behavioural patterns that govern species’ competitive outcomes (McMeans et al. 2020). As an example, in some cases, species may migrate (Holt and Fryxell 2011; Teitelbaum et al. 2015; Tavecchia et al. 2016; le Corre et al. 2020; Moorter et al. 2021) or hibernate (Campbell et al. 2008; Giraldo-Perez et al. 2016) during less productive seasons, opting out of competition when resources are scarce (Suski and Ridgway 2009). These behavioural strategies may counteract potential negative effects of periodic variation to some extent and thus promote coexistence by reducing competitive interactions during unfavourable times. More empirical research is needed to further understand these spatial and behavioural strategies, and seasonal trade-offs between competitive species, in order to understand their mechanistic role in maintaining biodiversity with climate change.

As nature abounds with temporal variation, grasping a better understanding of coexistence mechanisms in these fluctuating environments is now even more crucial with climate change. We have provided an analytical solution to begin understanding these effects. Our research has uncovered three competitive outcomes that could be found across a large range of natural periodicities and life history strategies. As species coexistence appears to be incredibly sensitive to periodic variation, climate change has the potential to drastically impact future competitive outcomes in the natural world.

## Acknowledgements

We would like to thank Cortland K. Griswold for his comments and suggestions that helped improve this manuscript. We would also like to thank the University of Guelph’s Elgin Card Scholarship in Terrestrial Ecology (Grant number [I5023]) awarded to AMS, and the NSERC Discovery Grant granted to KSM (Grant number [400353]) for funding this research.

## Code accessibility

All code to reproduce the above analyses and figures have been archived on Zenodo (version 1.0).

## Coexistence in Periodic Environments: Supplementary Material

### S1: A Linearization of the Periodic Lotka-Volterra Dynamics

If we consider any point on the phaseplane (*X1*-*X2*) due to the productive season that lasts from 0 to *τp* where *τ* is the period length, and *p* is the proportion of each period that is considered productive, then we know:

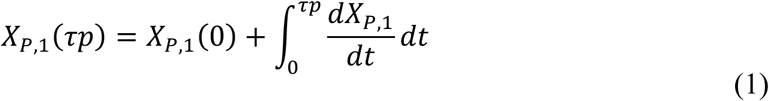

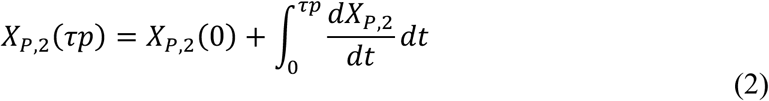

These dynamics follow the differential equation over the trajectory from 0 to *τp* starting at the values *XP,j*(0). We linearize the trajectory over 0 to *τp* by assuming the 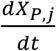 remains constant (e.g., we calculate the instantaneous 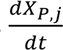 for a point in the phaseplane, say for time (0)). As this is now a constant, we can use the Fundamental Theorem of Calculus to solve for Equations (1) and (2) above yielding:

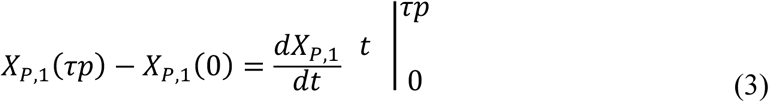

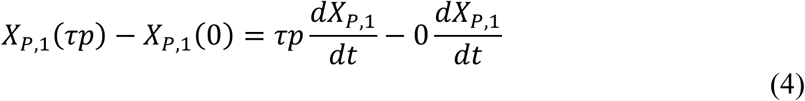

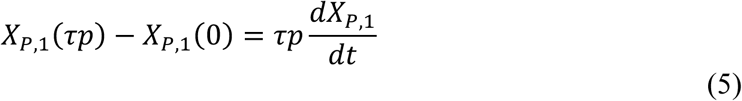

Similarly, we solve for the less productive season between *τp* and *τ*, giving us:

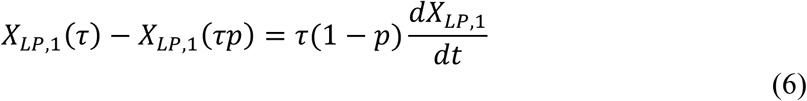

Thus, the linearization estimates the trajectory as a linear scale movement following the length of the time interval (either *τp*, or *τ*(1 – *p*)). We can take this simple estimate to estimate the within seasonal dynamics and use them to estimate the periodic 0-isoclines.

Given Equations (5) and (6), the *X1* isocline occurs when:

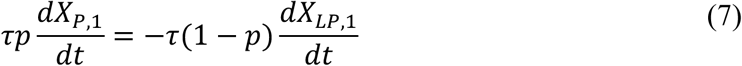

With this approximation, we substitute the productive and less productive Lotka-Volterra models into equation (7):

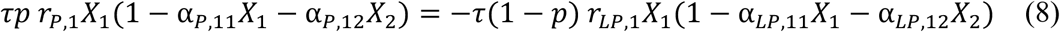

Noticing that *τ* cancels out, we can now solve for the isocline solution:

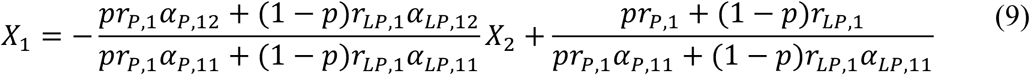

Similarly, we would perform the same steps to find the isocline solution for *X2*. This isocline approximation, for both species, allows us to determine coexistence criteria for seasonal environments.

### S2: Isocline Approximation and Numerical Simulation Comparison

**Fig. S2.1.**
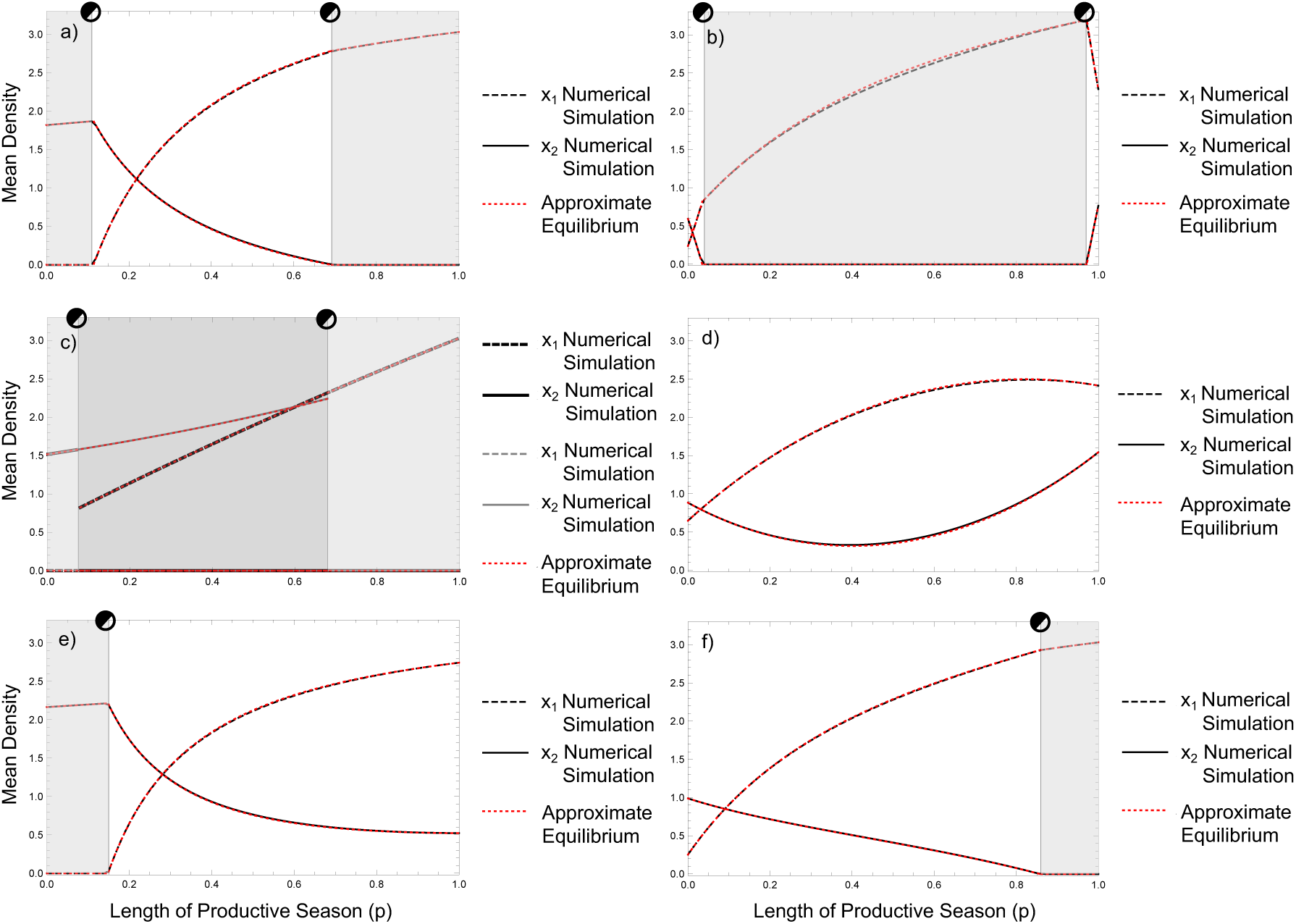
Comparison of Isocline Approximation against Numerical Simulations. White zone represents stable coexistence, light gray zone represents competitive exclusion, and dark gray zone represents contingent coexistence. a) Seasonally mediated coexistence. Parametric values: *αP,21*=0.35, *αP,12*=0.165, *αP,11*=0.33, *αP,22*=0.436, *rP,1*=1.7, *rP,2*=1.2, *αLP,21*=0.385, *αLP,12*=0,805, *αLP,11*=0.73, *αLP,22*=0.55, *rLP,1*=0.3, *rLP,2*=1. b) Seasonally mediated competitive exclusion. Parametric values: *αP,21*=0.29, *αP,12*=0.38, *αP,11*=0.31, *αP,22*=0.44, *rP,1*=4, *rP,2*=1.2, *αLP,21*=1.27, *αLP,12*=1.02, *αLP,11*=1,67, *αLP,22*=1.18 *rLP,1*=0.3, *rLP,2*=1. c) Seasonally mediated contingent coexistence. The equilibrium is unstable in the dark gray zone and multiple attractors exist between the two species. Parametric values: *αP,21*=0.548, *αP,12*=0.33, *αP,11*=0.33, *αP,22*=0.363, *rP,1*=1.7, *rP,2*=1.2, *αLP,21*=1.31, *αLP,12*=1.89, *αLP,11*=1.65, *αLP,22*=0.66, *rLP,1*=0.3, *rLP,2*=1. d) Counterintuitive mean density change. Parametric values: *αP,21*=0.2, *αP,12*=0.155, *αP,11*=0.315, *αP,22*=0.335, *rP,1*=1.7, *rP,2*=1.2, *αLP,21*=0.57, *αLP,12*=0.4, *αLP,11*=1, *αLP,22*=0.715, *rLP,1*=0.3, *rLP,2*=1. e) Mean density changes with competitive exclusion in the less productive season. Parametric values: *αP,21*=0.288, *αP,12*=0.15, *αP,11*=0.336, *αP,22*=0.402, *rP,1*=1.7, *rP,2*=1.2, *αLP,21*=0.354, *αLP,12*=0.75, *αLP,11*=0.666, *αLP,22*=0.462, *rLP,1*=0.3, *rLP,2*=1. f) Mean density changes with competitive exclusion in the productive season. Parametric values: *αP,21*=0.35, *αP,12*=0.165, *αP,11*=0.33, *αP,22*=0.44, *rP,1*=1.7, *rP,2*=1.2, *αLP,21*=0.28, *αLP,12*=0.825, *αLP,11*=0.73, *αLP,22*=0.94, *rLP,1*=0.3, *rLP,2*=1.

**Fig. S2.2.**
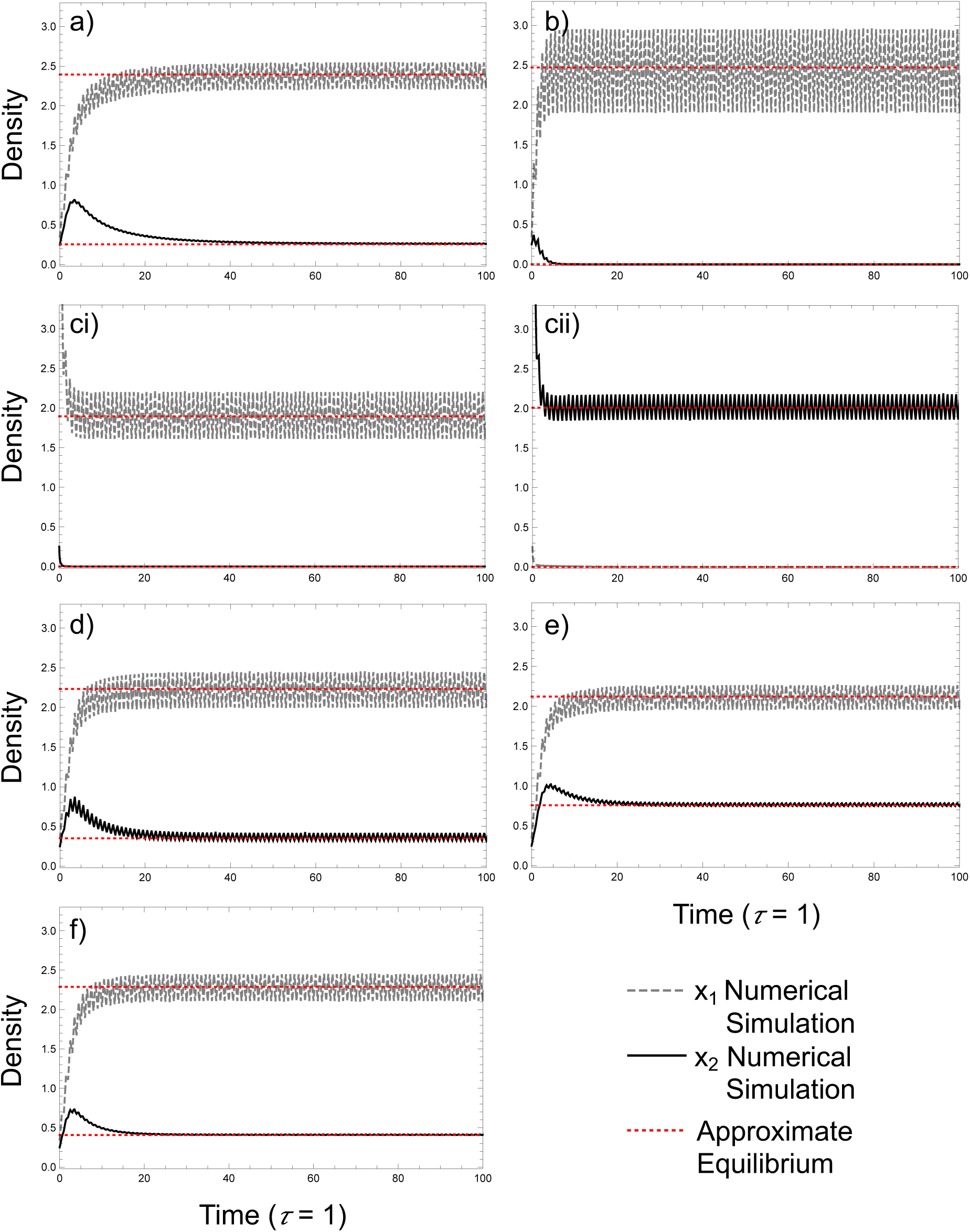
Time Series at *p* = 0.5, and *τ* = 1 for the comparison of isocline approximation against numerical simulations. a) Seasonally mediated coexistence. Parametric values: *αP,21*=0.35, *αP,12*=0.165, *αP,11*=0.33, *αP,22*=0.436, *rP,1*=1.7, *rP,2*=1.2, *αLP,21*=0.385, *αLP,12*=0,805, *αLP,11*=0.73, *αLP,22*=0.55, *rLP,1*=0.3, *rLP,2*=1. b) Seasonally mediated competitive exclusion. Parametric values: *αP,21*=0.29, *αP,12*=0.38, *αP,11*=0.31, *αP,22*=0.44, *rP,1*=4, *rP,2*=1.2, *αLP,21*=1.27, *αLP,12*=1.02, *αLP,11*=1,67, *αLP,22*=1.18 *rLP,1*=0.3, *rLP,2*=1. ci) Seasonally mediated contingent coexistence. *X1* competitively excluded *X2*. Parametric values: *αP,21*=0.548, *αP,12*=0.33, *αP,11*=0.33, *αP,22*=0.363, *rP,1*=1.7, *rP,2*=1.2, *αLP,21*=1.31, *αLP,12*=1.89, *αLP,11*=1.65, *αLP,22*=0.66, *rLP,1*=0.3, *rLP,2*=1. cii) Seasonally mediated contingent coexistence. *X2* competitively excluded *X1*. Parametric values: *αP,21*=0.548, *αP,12*=0.33, *αP,11*=0.33, *αP,22*=0.363, *rP,1*=1.7, *rP,2*=1.2, *αLP,21*=1.31, *αLP,12*=1.89, *αLP,11*=1.65, *αLP,22*=0.66, *rLP,1*=0.3, *rLP,2*=1. d) Counterintuitive mean density change. Parametric values: *αP,21*=0.2, *αP,12*=0.155, *αP,11*=0.315, *αP,22*=0.335, *rP,1*=1.7, *rP,2*=1.2, *αLP,21*=0.57, *αLP,12*=0.4, *αLP,11*=1, *αLP,22*=0.715, *rLP,1*=0.3, *rLP,2*=1. e) Mean density changes with competitive exclusion in the less productive season. Parametric values: *αP,21*=0.288, *αP,12*=0.15, *αP,11*=0.336, *αP,22*=0.402, *rP,1*=1.7, *rP,2*=1.2, *αLP,21*=0.354, *αLP,12*=0.75, *αLP,11*=0.666, *αLP,22*=0.462, *rLP,1*=0.3, *rLP,2*=1. f) Mean density changes with competitive exclusion in the productive season. Parametric values: *αP,21*=0.35, *αP,12*=0.165, *αP,11*=0.33, *αP,22*=0.44, *rP,1*=1.7, *rP,2*=1.2, *αLP,21*=0.28, *αLP,12*=0.825, *αLP,11*=0.73, *αLP,22*=0.94, *rLP,1*=0.3, *rLP,2*=1.

### S3: Tracking Isocline Approximation across Productivity Length for all Seasonally

**Fig. S3.1.**
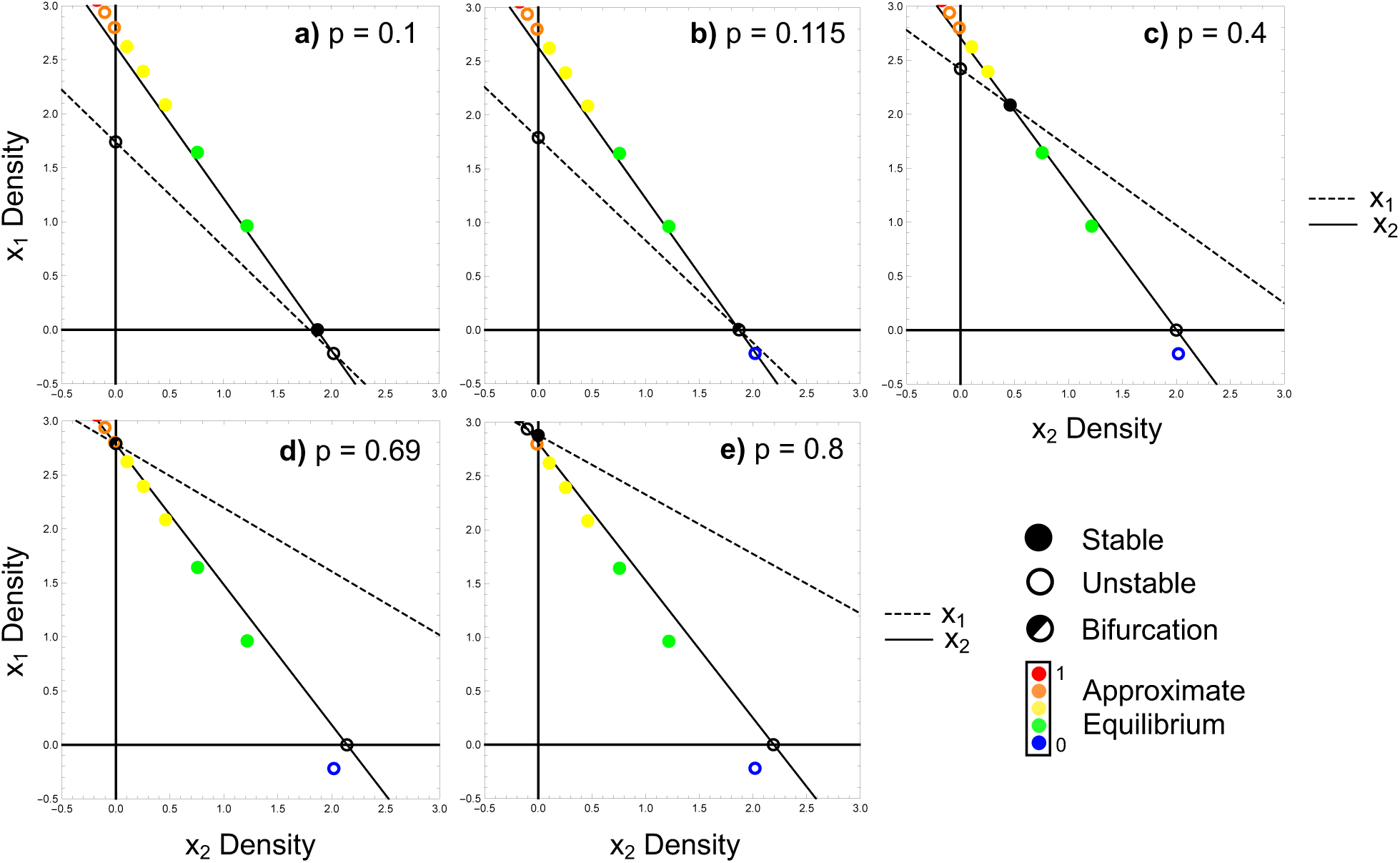
Seasonally mediated coexistence isocline approximation. Isocline approximation tracking the interior equilibrium as the productive season increases in length from a) *p*=0.1; competitive exclusion of species 1, to b) *p*=0.115; transcritical bifurcation, to c) *p*=0.4; stable coexistence, to d) *p*=0.69; transcritical bifurcation, to e) *p*=0.8; competitive exclusion of species 2. Parametric values: *αP,21*=0.35, *αP,12*=0.165, *αP,11*=0.33, *αP,22*=0.436, *rP,1*=1.7, *rP,2*=1.2, *αLP,21*=0.385, *αLP,12*=0,805, *αLP,11*=0.73, *αLP,22*=0.55, *rLP,1*=0.3, *rLP,2*=1.

**Fig. S3.2.**
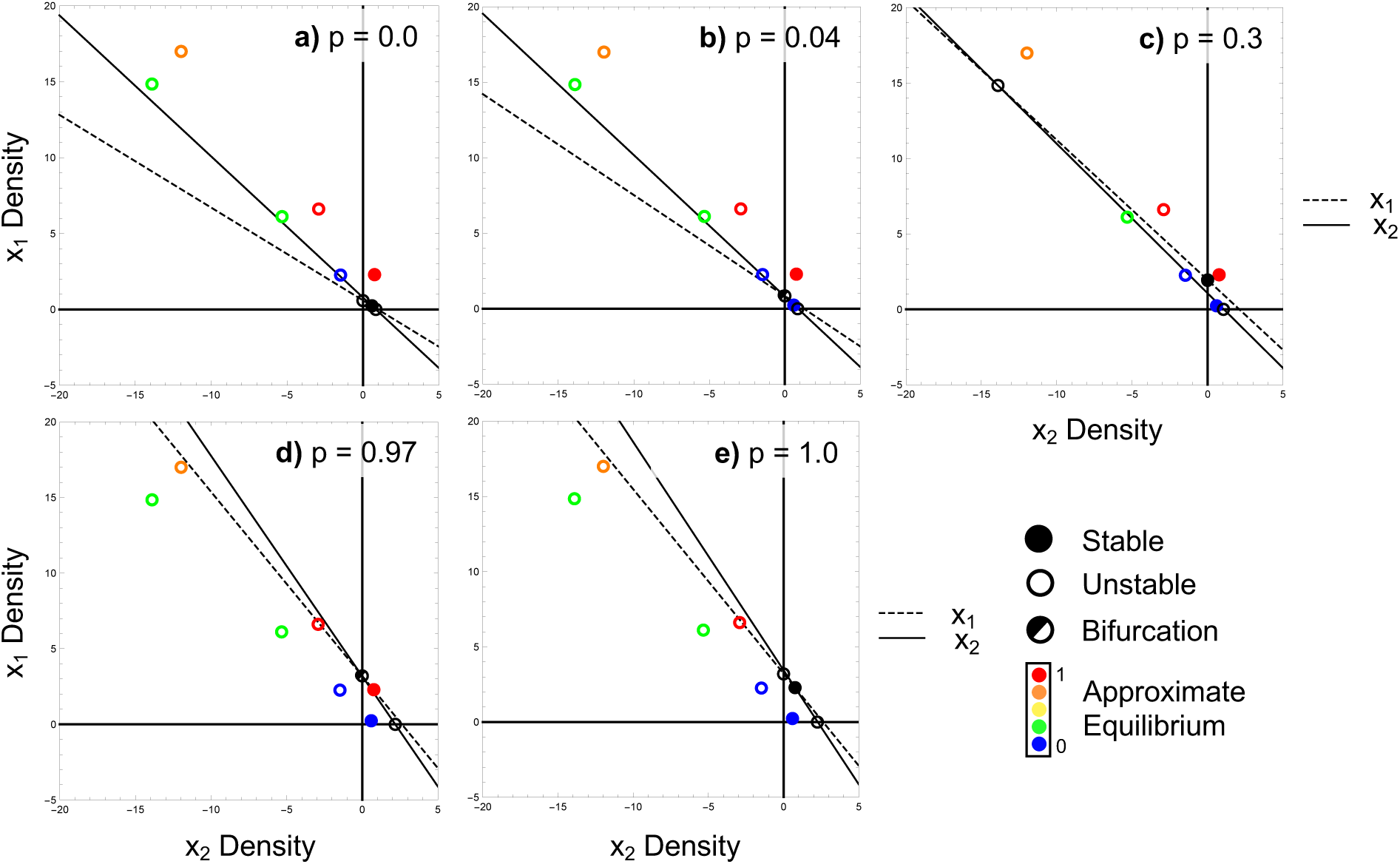
Seasonally mediated competitive exclusion isocline approximation. Isocline approximation tracking the interior equilibrium as the productive season increases in length from a) *p*=0.0; stable coexistence, to b) *p*=0.04; transcritical bifurcation, to c) *p*=0.3; competitive exclusion of species 2, to d) *p*=0.97; transcritical bifurcation, to e) *p*=1.0; stable coexistence. Parametric values: *αP,21*=0.29, *αP,12*=0.38, *αP,11*=0.31, *αP,22*=0.44, *rP,1*=4, *rP,2*=1.2, *αLP,21*=1.27, *αLP,12*=1.02, *αLP,11*=1,67, *αLP,22*=1.18 *rLP,1*=0.3, *rLP,2*=1.

**Fig. S3.3.**
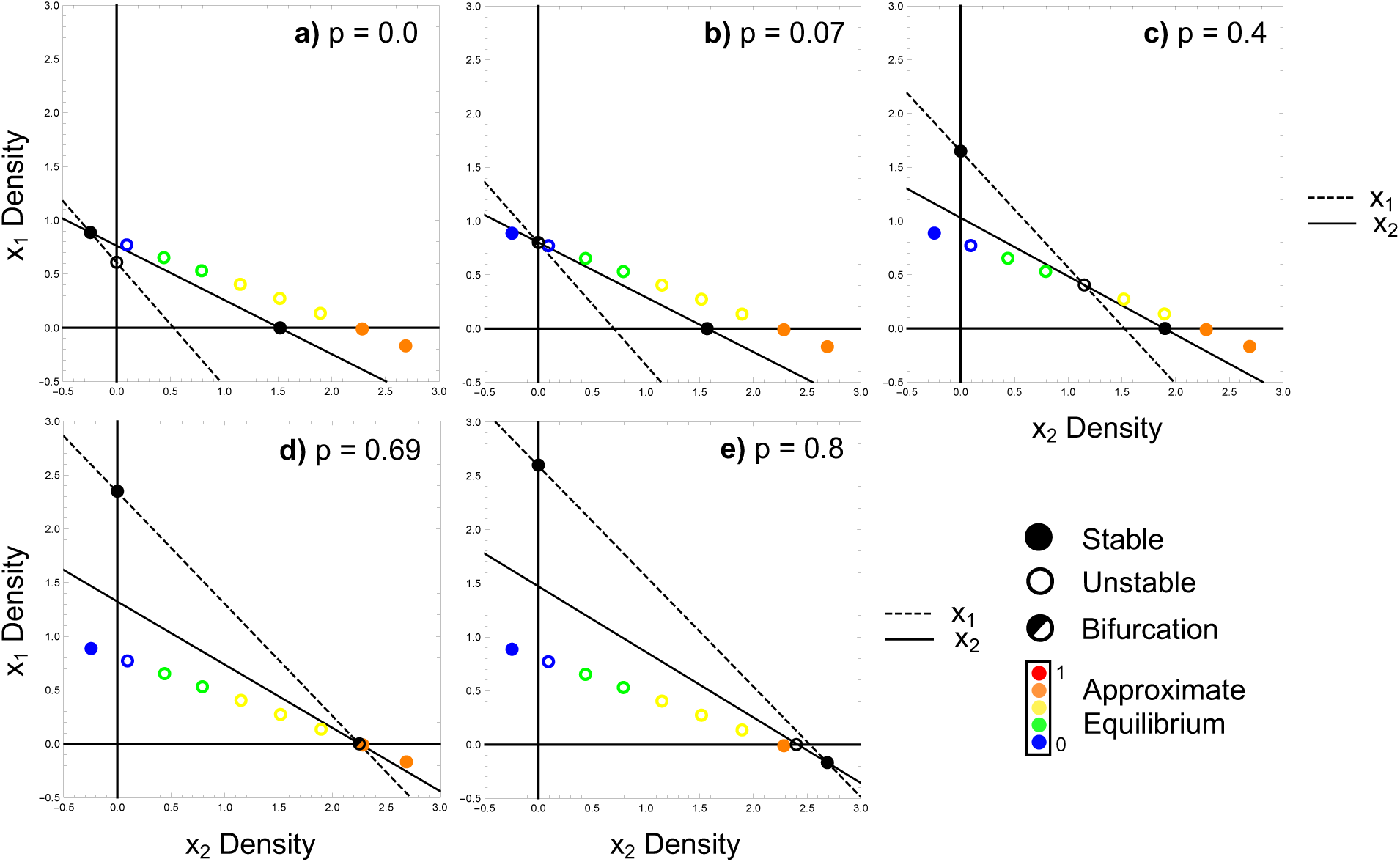
Seasonally mediated contingent coexistence isocline approximation. Isocline approximation tracking the interior equilibrium as the productive season increases in length from a) *p*=0.0; competitive exclusion of species 1, to b) *p*=0.07; transcritical bifurcation, to c) *p*=0.4; contingent coexistence, to d) *p*=0.69; transcritical bifurcation, to e) *p*=0.8; competitive exclusion of species 2. Parametric values: *αP,21*=0.548, *αP,12*=0.33, *αP,11*=0.33, *αP,22*=0.363, *rP,1*=1.7, *rP,2*=1.2, *αLP,21*=1.31, *αLP,12*=1.89, *αLP,11*=1.65, *αLP,22*=0.66, *rLP,1*=0.3, *rLP,2*=1.

### S4: Accuracy of the Linear Isocline Approximation

Our results hold for a broad range of period lengths but clearly depend on specific parameterizations that alter the timescale or the effects of timescale. Recall that our approximation operates by assuming a zero population growth rate exists in the phaseplane where linearization’s of the Lotka-Volterra equations cancel out over the high growth and low growth periods (i.e., linearization of the high growth for species *X1* and species *X2* are both equal and opposite in sign to the low growth linearization). Given this assumption, we can immediately ask when we expect nonlinear dynamics to dominate and potentially threaten the validity of the linear assumption. First, the longer the period of the forced parameters (*τ*), then the longer the time the dynamics have to fall off the linear assumption. Further, the larger the growth rates (*r*), the larger the potential for nonlinear dynamics even with smaller periods (*τ*). As a result, we can say the larger the product *rτ,* the more likely our assumption of linearity is threatened.

Effectively, *τr* sets the relative pace of the seasonal dynamics.

Recognizing that linear assumptions will lose accuracy when dynamics become nonlinear with longer periods and fast growth rates (i.e., there is more time for dynamics to become nonlinear), we first compare the dynamics (set by the original growth rates in the manuscript) with the approximation’s accuracy when the period length (*τ*) is increased. Next, we slow the dynamics by dividing all species’ growth rates (*r*) by 10-units and compare the approximation’s accuracy with these dynamics when the period length increases. Our results below show that indeed our approximation can fall off, but even here for large, combined values of *rτ*, the approximation remains a reasonable qualitative predictor of steady state behaviour for seasonally-mediated coexistence and contingent coexistent (S4.1 and S4.3 respectively). We note that the seasonally mediated competitive exclusion is not as robust (S4.2).

As seen in Fig. 4a, the analytical approximation still accurately predicts the numerically generated transcritical bifurcation at *p* = 0.11 (mean densities transition between competitive exclusion of species 1 to coexistence of both species), even when the period length (*τ*), reaches 100 time-units. This is due to the large difference between the productive and less-productive growth rates of species 1. However, the approximation begins to inaccurately predict where the second numerically generated transcritical bifurcation will occur (*p* = 0.69; mean density transition between coexistence and competitive exclusion of species 2) when *τ* = 5 time-units (Fig. S4.11b). With the original growth rates used in the manuscript, when the period length is small (*τ* = 1), the seasonal dynamics are relatively linear, and the approximation accurately predict the true mean densities from the numerical simulation (Figs. S4.11a and S4.12a).

However, as we increase the period length to 5 time-units, the dynamics become more nonlinear as, with enough time, species’ densities reach their seasonal equilibrium and spend more time at these fixed points (Fig. S4.12b). Here, the numerically generated mean densities will be skewed closer towards this equilibrium, while the linear approximation fails to capture this (Fig. S4.11b). When all growth rates are slowed down (all *r*’s divided by 10 units), the seasonal dynamics become more linear, and the approximation is more accurately able to track the mean densities even when the period length (*τ*) is increased from 1 (Figs S4.11c and S4.12c) to 5 time-units (Figs S4.11d and S4.12d).

In Fig. 5a, the approximation still predicts seasonally mediated competitive exclusion, even when species coexist at very large period lengths (*τ* > 33 time units) across all *p*-values.

However, when *τ* is increased from 1 to 10 time-units with the original growth rates, the dynamics become more nonlinear (Figs. S4.22a and S4.22b respectively) and the approximation begins to fail at tracking the numerically-generated mean densities throughout the entire range of *p*-values (Figs. S4.21a and S4.21b respectively), though it still captures the qualitative behavior and presence of the bifurcations. When growth rates are smaller, the seasonal dynamics are now more linear (Figs. S4.22c and FigsS4.22d), and the approximation more accurately tracks the numerically generated mean densities as *τ* increases from 1 to 10 time-units (Figs. S4.22c and FigsS4.22d respectively).

For seasonally mediated contingent coexistence, as the period length (*τ*) increases, the region of contingent coexistence from the numerically-generated results first broadens and then shrinks (Fig. 6a). With the original growth rates, as the period length is increased from 1 to 20 time- units, the seasonal dynamics become more nonlinear (Figs. S4.32a and S4.32b respectively), and the approximation is unable to track the true bifurcations based on our numerical results (Figs. S4.31a and S4.31b respectively). When the growth rates are a tenth of their original size (all *r*’s divided by 10 units), the dynamics slow down and are now more linear as *τ* is increased (Figs. S4.32c and S4.32d), allowing the approximation to more accurately track the numerically- generated mean densities over the entire range of *p*-values (Figs. S4.31c and S4.31d).

### S4.1 Approximation Accuracy of Seasonally Mediated Coexistence

**Fig. S4.11.**
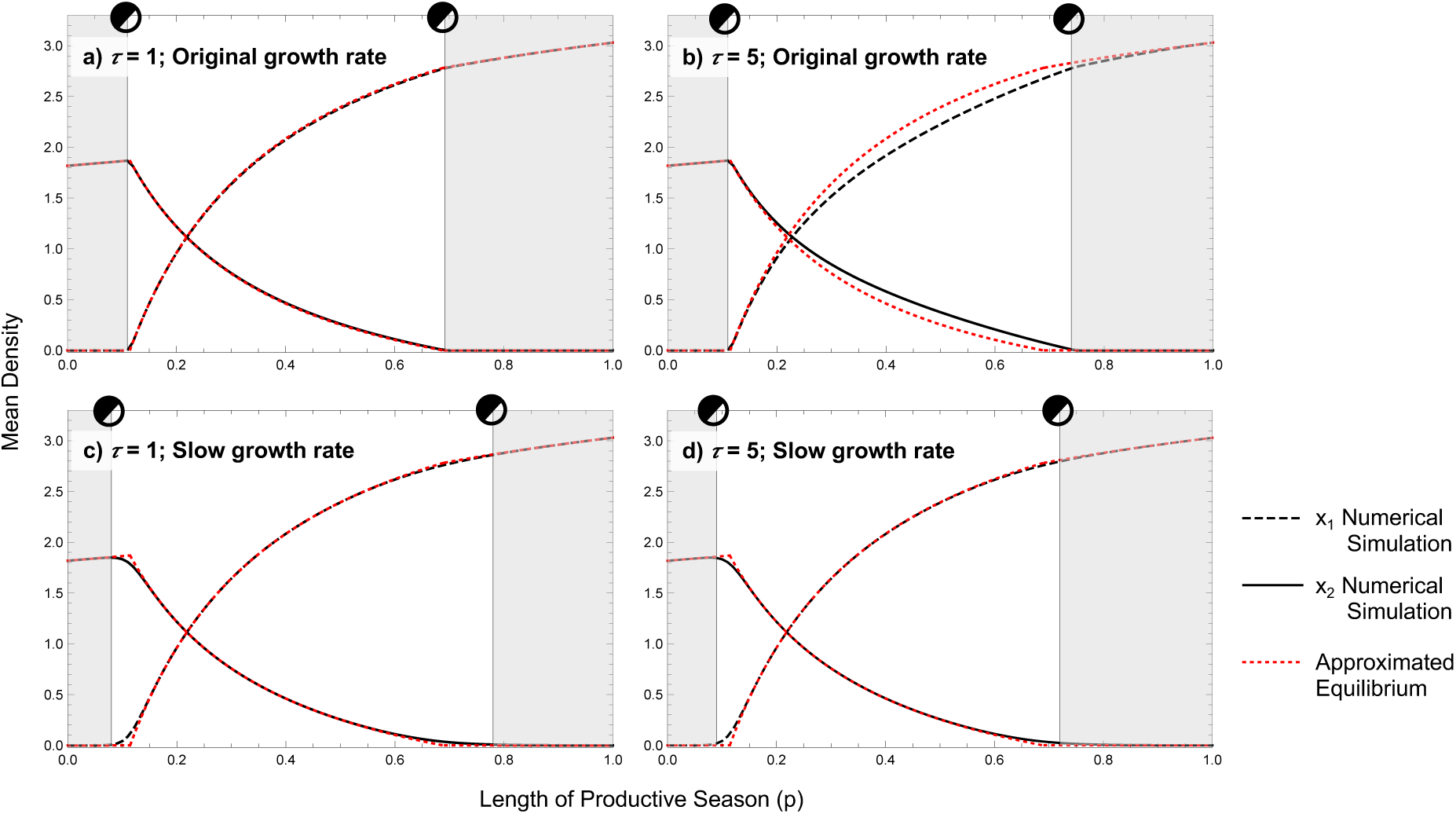
Isocline approximation – mean density plots over the length of the productive season (*p*) for seasonally-mediated coexistence. White zones represent coexistence and light grey zones represent competitive exclusion. a) and b) original growth rates (those used throughout manuscript: *rP1* = 1.7, *rP2* = 1.2, *rLP1* = 1, *rLP2* = 0.3) when the period length *τ* = 1 and 5 time- units respectively. c) and d) all growth rates have been divided by 10 units (*rP1* = 0.17, *rP2* = 0.12, *rLP1* = 0.1, *rLP2* = 0.03) when the period length = 1 and 5 time-units respectively. Parametric values: *αP,21*=0.35, *αP,12*=0.165, *αP,11*=0.33, *αP,22*=0.436, *αLP,21*=0.385, *αLP,12*=0,805, *αLP,11*=0.73, *αLP,22*=0.55.

**Fig. S4.12.**
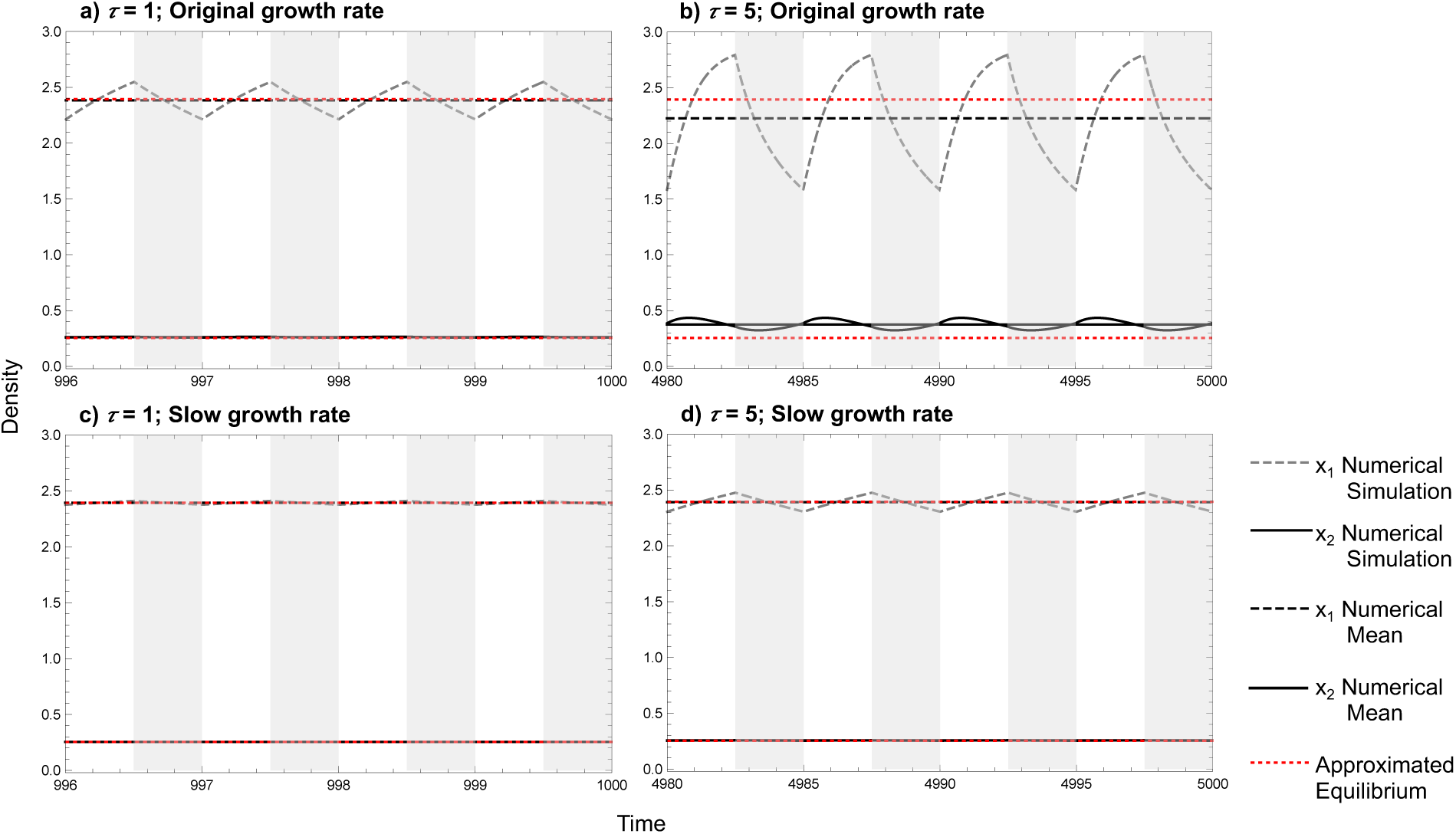
Isocline approximation – time series for seasonally-mediated coexistence. Length of productive season (*p*) = 0.5 represented by white zones, and light grey zones represent the less-productive season. a) and b) original growth rates (those used throughout manuscript: *rP1* = 1.7, *rP2* = 1.2, *rLP1* = 1, *rLP2* = 0.3) when the period length *τ* = 1 and 5 time-units respectively. c) and d) all growth rates have been divided by 10 units (*rP1* = 0.17, *rP2* = 0.12, *rLP1* = 0.1, *rLP2* = 0.03) when the period length = 1 and 5 time-units respectively. Parametric values: *αP,21*=0.35, *αP,12*=0.165, *αP,11*=0.33, *αP,22*=0.436, *αLP,21*=0.385, *αLP,12*=0,805, *αLP,11*=0.73, *αLP,22*=0.55.

### S4.2 Approximation Accuracy of Seasonally Mediated Competitive Exclusion

**Fig. S4.21.**
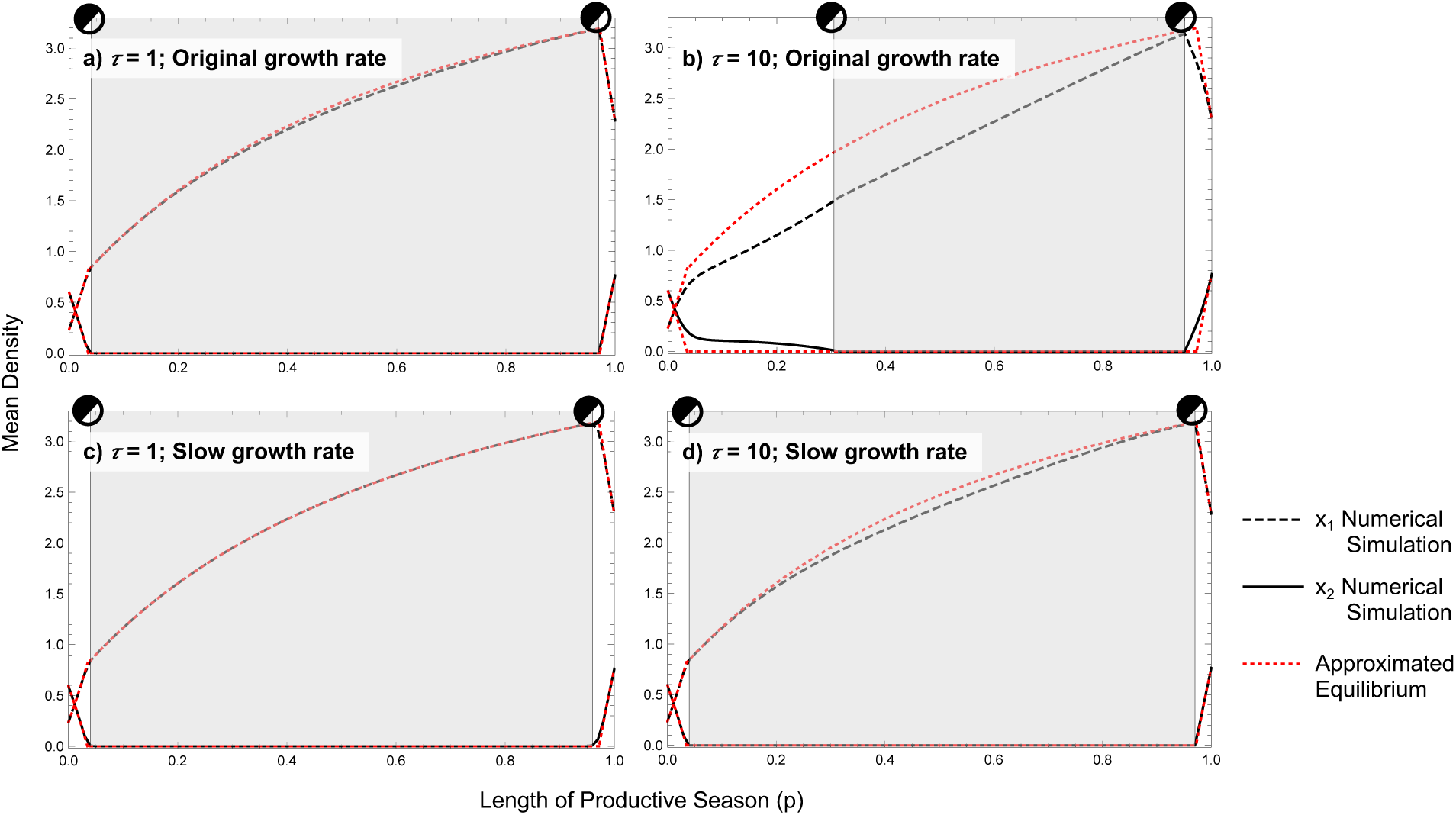
Isocline approximation – mean density plots over the length of the productive season (*p*) for seasonally-mediated competitive exclusion. White zones represent coexistence and light grey zones represent competitive exclusion. a) and b) original growth rates (those used throughout manuscript: *rP1* = 4, *rP2* = 1.2, *rLP1* = 1, *rLP2* = 0.3) when the period length *τ* = 1 and 10 time-units respectively. c) and d) all growth rates have been divided by 10 units (*rP1* = 0.4, *rP2* = 0.12, *rLP1* = 0.1, *rLP2* = 0.03) when the period length = 1 and 10 time-units respectively. Parametric values: *αP,21*=0.29, *αP,12*=0.38, *αP,11*=0.31, *αP,22*=0.44, *αLP,21*=1.27, *αLP,12*=1.02, *αLP,11*=1,67, *αLP,22*=1.18.

**Fig. S4.22.**
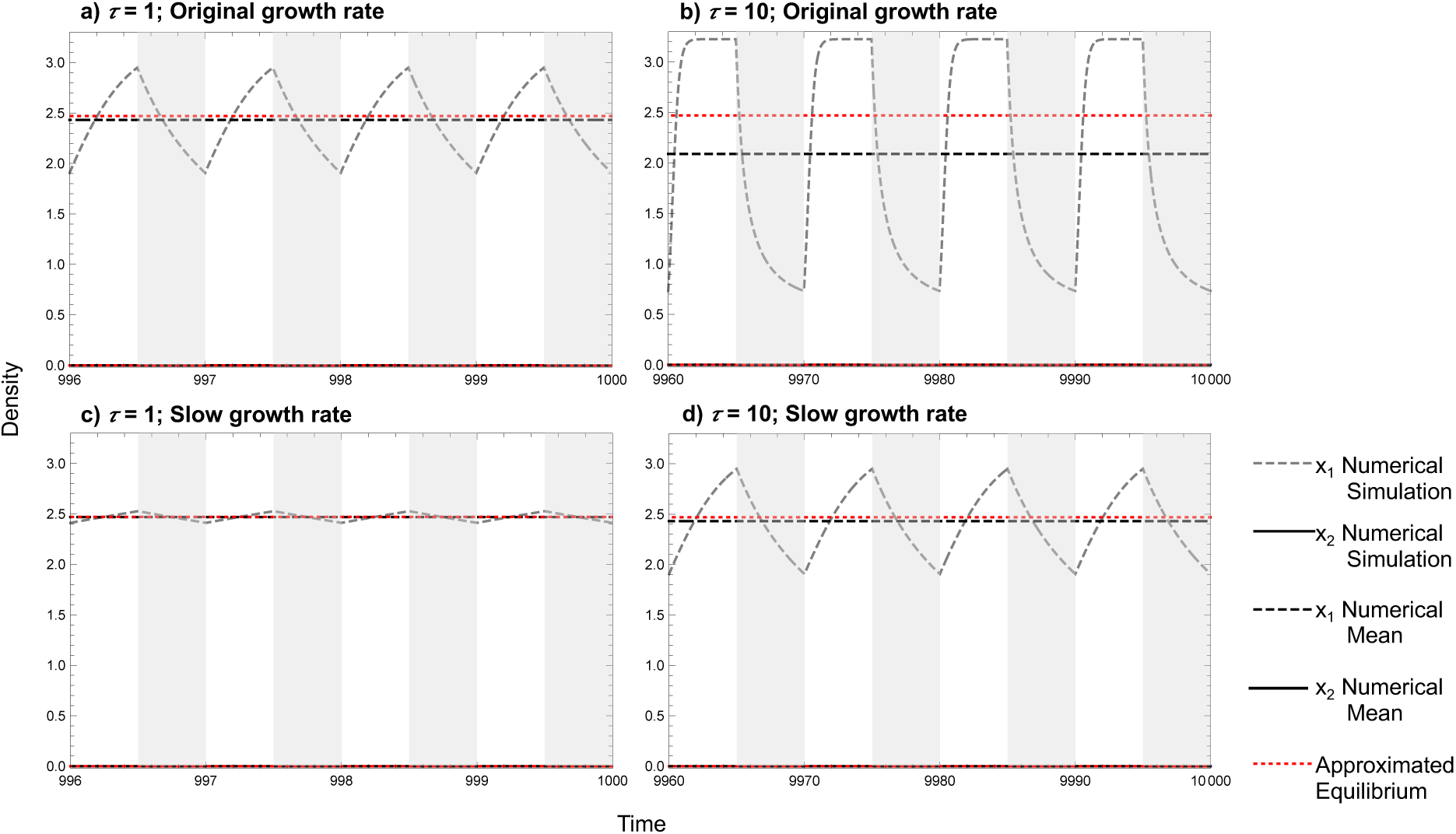
Isocline approximation – time series for seasonally-mediated competitive exclusion. Length of productive season (*p*) = 0.5 represented by white zones, and light grey zones represent the less-productive season. a) and b) original growth rates (those used throughout manuscript: *rP1* = 4, *rP2* = 1.2, *rLP1* = 1, *rLP2* = 0.3) when the period length *τ* = 1 and 10 time-units respectively. c) and d) all growth rates have been divided by 10 units (*rP1* = 0.4, *rP2* = 0.12, *rLP1* = 0.1, *rLP2* = 0.03) when the period length = 1 and 10 time-units respectively. Parametric values: *αP,21*=0.29, *αP,12*=0.38, *αP,11*=0.31, *αP,22*=0.44, *αLP,21*=1.27, *αLP,12*=1.02, *αLP,11*=1,67, *αLP,22*=1.18.

### S4.3 Approximation Accuracy of Seasonally Mediated Contingent Coexistence

**Fig. S4.31.**
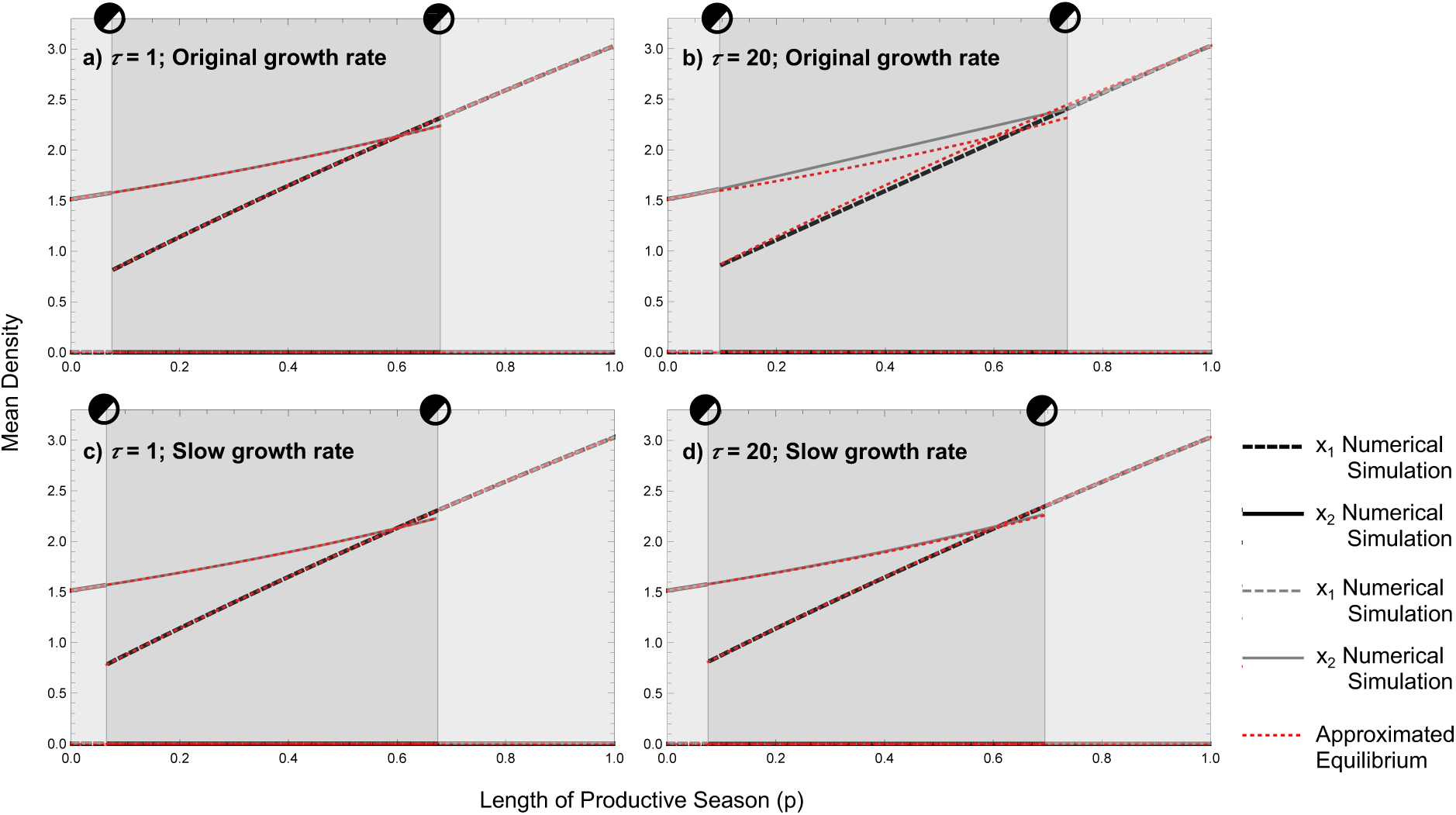
Isocline approximation – mean density plots over the length of the productive season (*p*) for seasonally-mediated contingent coexistence. Light grey zones represent competitive exclusion and grey zones represent contingent coexistence. a) and b) original growth rates (those used throughout manuscript: *rP1* = 1.7, *rP2* = 1.2, *rLP1* = 1, *rLP2* = 0.3) when the period length *τ* = 1 and 20 time-units respectively. c) and d) all growth rates have been divided by 10 units (*rP1* = 0.17, *rP2* = 0.12, *rLP1* = 0.1, *rLP2* = 0.03) when the period length = 1 and 20 time-units respectively. Parametric values: *αP,21*=0.548, *αP,12*=0.33, *αP,11*=0.33, *αP,22*=0.363, *αLP,21*=1.31, *αLP,12*=1.89, *αLP,11*=1.65, *αLP,22*=0.66.

**Fig. S4.32.**
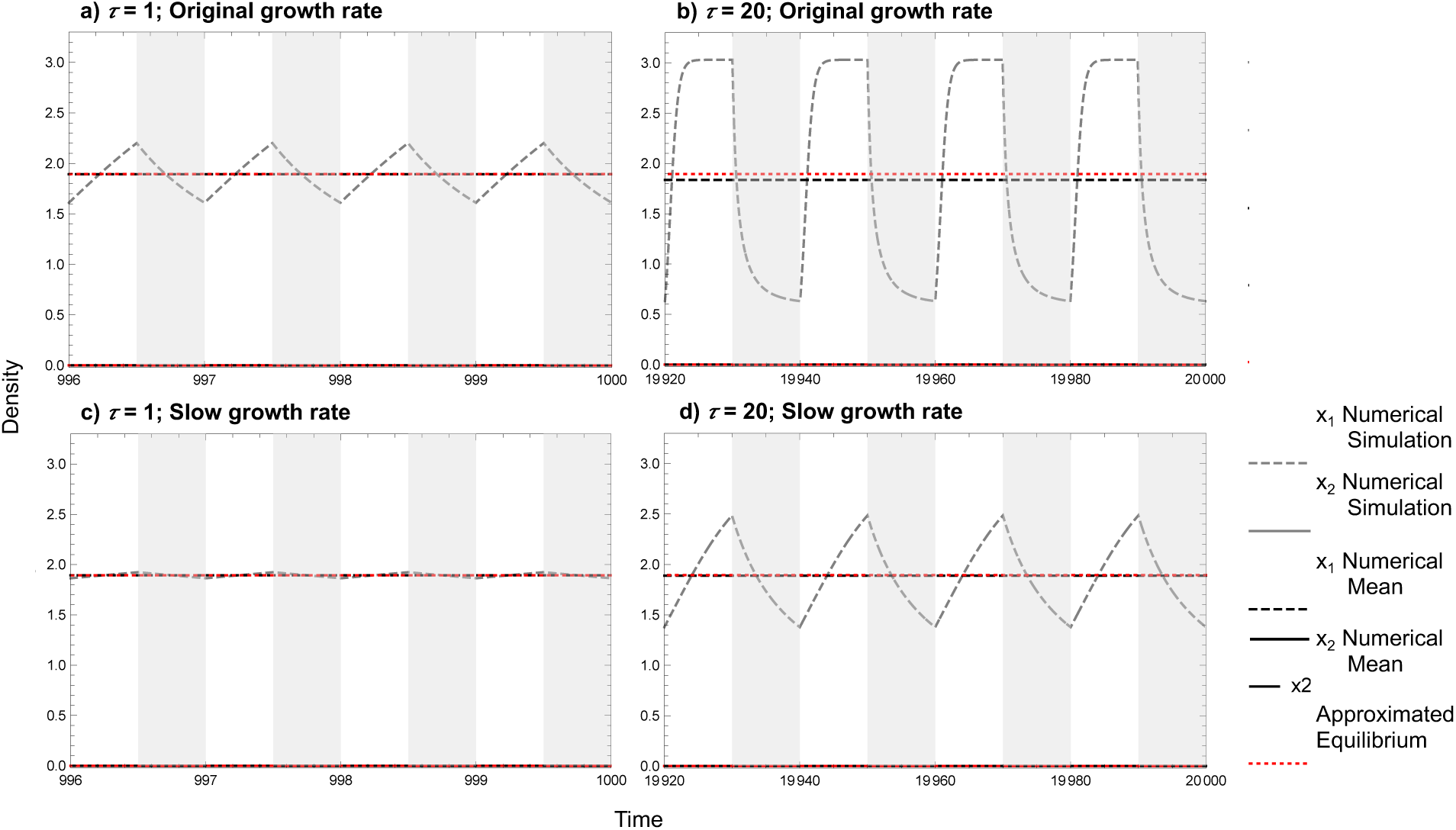
Isocline approximation – time series for seasonally-mediated contingent coexistence. Length of productive season (*p*) = 0.5 represented by white zones, and light grey zones represent the less-productive season. a) and b) original growth rates (those used throughout manuscript: *rP1* = 1.7, *rP2* = 1.2, *rLP1* = 1, *rLP2* = 0.3) when the period length *τ* = 1 and 20 time-units respectively. c) and d) all growth rates have been divided by 10 units (*rP1* = 0.17, *rP2* = 0.12, *rLP1* = 0.1, *rLP2* = 0.03) when the period length = 1 and 20 time-units respectively. Parametric values: *αP,21*=0.548, *αP,12*=0.33, *αP,11*=0.33, *αP,22*=0.363, *αLP,21*=1.31, *αLP,12*=1.89, *αLP,11*=1.65, *αLP,22*=0.66.

